# Electron transfer parameters for Methemoglobin formation in mutant Hemoglobin α-chains

**DOI:** 10.1101/2021.03.28.437393

**Authors:** Vaibhav A. Dixit, Jochen Blumberger, Shivam Kumar Vyas

## Abstract

Hemoglobin mediated transport of dioxygen (O_2_) critically depends on the stability of the reduced (Fe^2+^) form of the Heme cofactors. Some protein mutations stabilize oxidized (Fe^3+^) state (Methemoglobin, Hb M) causing methemoglobinemia and can be lethal above 30 %. Majority of the analyses of factors influencing Hb oxidation are retrospective and give insights only for inner sphere mutations of Heme (His58, His87). Herein, we report the first all atom MD simulations on redox states and calculations of the Marcus ET parameters for the α-chain Hb oxidation and reduction rates for Hb M. The Hb (wild type), and most of the studied α-chain variants maintain globin structure except the Hb M Iwate (H87Y). Using linear response approximation we calculated average energy gaps (<ΔE>), total (λ), protein (λ_prot_), solvent (λ_solv_) reorganization energies, and redox potentials (*E*°), and oxidation free energies (ΔG°). The total λ ranges from 0.685 – 0.730 eV in agreement with literature on Hb and similar Heme proteins. The mutants forming Hb M tend to lower the *E*° and thus stabilize the oxidized (Fe^3+^) state (e.g. the Hb Miyagi variant with K61E mutation). Solvent reorganization (λ_solv_ 73 – 96 %) makes major contributions to λ, while protein reorganization (λ_prot_) accounts for 27 – 30 % except for the Miyagi and J-Buda variants (λ_prot_ ∼ 4 %). Analysis of Heme-solvent H-bonding interactions among variants provide insights into the role of Lys61 residue in stabilizing Fe^2+^ state and ET parameters. The ET parameters provide valuable insights into the Hb oxidation to Hb M in agreement with the experimental data. Thus our methodology explains the effect of mutations on the structure, stability and Hb oxidation, and has potential for the prediction of methemoglobinemia.

## 1. Introduction

Hemoglobin variants are wide spread in the Human population as seen in the HbVar database (http://globin.cse.psu.edu/hbvar/menu.html). These variants lead to two broad classes of disorders: 1) Globin synthesis and assembly disorders known as Thalassemias and 2) Globin structure disorders (Methemoglobinemia, Sickle cell anemia, and disorders caused by unstable, and altered dioxygen (O_2_) affinity Hb variants).^1^ Hb variant characterization, biochemical properties and associated clinical manifestations have been reviewed recently.^2,3^ Hb is normally present in the reduced (Fe^2+^, 99%) state and effectively transports O_2_ between lungs and tissues. In red blood cells (RBCs), the concentration of the oxidized Hb, Methemoglobin (metHb, also known as Hb M), is kept below 1 % by enzymatic (Cytochrome b5, and NADPH dependent Flavin reductase) and non-enzymatic (e.g. ascorbic acid) reductive pathways.^4^ Higher levels (∼ 10 %) can lead to cyanosis, reduced O_2_ delivery and level above 30 % can be lethal. This condition, known as Methemoglobinemia, can be induced by drugs (chloroquine/hydroxychloroquine), inherited defects in Cytochrome b5 (or its reductase), or inheritance of an Hb variant characterized by an increase in the stability of oxidized (Ferric, Fe^3+^) or a decrease in the stability of the (ferrous, Fe^2+^) states.^5–10^ Methemoglobinemia has been reported in many COVID-19 patients,^11–13^ thus potentially complicating the assessment of blood O_2_ saturation levels in respiratory distress. Incidence of COVID-19 in patients with sickle cell disease (characterized by Hb S) was found to be 85 %, but prevalence of other Hb variants in COVID-19 patients remains unexplored.^14^

A large body of information and knowledge is available in the form of more than 1820 known variants, > 600 human Hb structures in the PDB, a large number of biochemical, genetic, spectroelectrochemical, and modeling studies. The structure and function of Hb and impact of mutations on O_2_ affinity has been reviewed in the past.^15–17^ The Hb globin structure has eight helices that maintain a hydrophobic environment around the Heme and stabilize Fe^2+^ state to ensure reversible O_2_ binding. Structural changes that destabilize the goblin or expose Heme to solvent enhance oxidation rates and Hb M formation. Robust clinical and biochemical tests (HPLC, Electrophoretic, MALDI-TOF and PCR), are available for the characterization of different variants, for example sickle cell Hb (Hb S), and thalassemic forms (Hb F),^3^ but functional characterization of these variants in terms of O_2_ affinity or Hb M content, oxidation, and reduction rates are extremely rare.^18^

Thus the in vivo rates for Hb oxidation to Hb M and reduction of Hb M back to Fe^2+^ state for many Hb variants remains unknown. These unknowns can potentially lead to failures in diagnosis, delays in the treatment, overtreatment and even cause treatment failures with standard therapy like methylene blue.^9,19^ Additionally, due to the lack of knowledge of factors like optimum reductant concentrations, and O_2_ pressures, Hb M content can go unrecognized and reach toxic levels causing cyanosis and even fatality. Thus reliable estimation of the relative Hb oxidation and Hb M reduction rates for different Hb variants is urgently required.

Marcus theory of electron transfer (ET) and its variants have been successfully applied in the areas of material science, small molecule charge-transfer complexes, redox heme and non-heme proteins involved in respiratory and mitochondrial ET chains.^20^ Since Hb is a tetramer with two identical α and β chains which contain identical Iron-porphyrin (heme) co-factor, experimental measurements of enzymatic and non-enzymatic Hb reduction rates for individual subunits and applications of ET theories has proved challenging. McLendon et al., have used reconstituted [Zn, Fe] hybrid Hb dimers in which Heme-Fe in one of the chains has been replaced with znic.^21^ These hybrid Hbs show a larger free energy change (ΔG°) associated with the reduction (Equation 1) compared to the wild type Hb. Using Marcus theory and assuming a constant reorganization energy of (λ = 0.9 V) allowed the prediction of the ET rates (*k*_ET_) for the wild and hybrid Hb. The wild type *k*_ET_ = 1.6 s^-1^ was predicted in close agreement with experimental value of 1 and 2 s^-1^ at pH = 7 and 6 respectively. ET rates from the photoinduced excited state of hybrid Hb were about 1500 fold faster due to a larger ΔG°. Based on the binding constant (k_b_) and observed *k*_ET_, the authors concluded that formation of a specific biomolecular complex between Cyt b5 and Hb may not be necessary and ET takes places via a simple biomolecular collision mechanism.

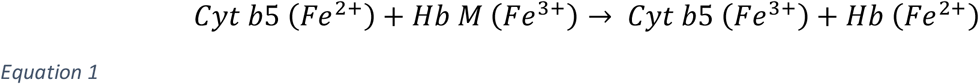

Earlier studies by Mauk et al., had found contrasting spectroscopic evidence with quantitative formation of 1:1 Cyt b5:Hb complexes.^22^ Extending this work they proposed a model for the Cyt b5:Hb complex in which the Heme edges are 8 Å apart while the Fe-Fe distance in the complex was 16 Å.^23^ Hb residues (Lys 54, 56, 60, 61, 90, heme propionates) and Cyt b5 residues (Glu 43, 44, 48, ASP 60, and heme propionates) were postulated to be involved in complex formation. Later Hoffman et al., used similar Hb hybrids and determined the ET rates for individual Hb subunits.^24^ The α subunit showed 4 fold higher binding affinity and 2 fold faster ET rates compared to the β subunits. These findings were similar to earlier reports of the ET between Hb and inorganic complexes.^25^ Brittan et al., performed Brownian dynamics and electrostatics calculations to reassess the structural requirements for efficient electron transfer between Cyt b5 and embryonic Hb.^26^ Although there was a descent agreement with Mauk et al,^22^ regarding the identity of residues involved in the complex formation, the ET rates were overestimated for the native Hb and were similar to the hybrid Hb. None of these studies have performed all atom MD simulations of the redox states to estimate Marcus parameters and ET rates.

The experimental redox potentials (*E*°) for Hb and Hb subunits depend on the details of the experimental method (direct electrochemistry, or spectroelectrochemistry), nature of the electrodes and its interactions with Hb, and conditions e.g. presence of O_2_, pH (Bohr effect), ionic strength, redox mediator, and temperature. Literature values range from -0.172 to + 0.050 V. In studies with the Hb tetramer, the identity of the subunits undergoing the redox changes are uncertain. Earlier Mateescu et al., have performed cyclic voltammetry experiments to determine the Hb *E*°s under acidic/aerobic and neutral/anaerobic conditions which are known to stabilize the T (tense) and R (relaxed) states respectively.^27^ The authors assigned *E*° values of -0.040 and -0.165 V respectively to the R and T states of Hb. Guiles et al, have measured the *E*° of Hb, Cyt b5 and used Marcus theory to predict reorganization energies (λ) and homogenous and heterogenous ET rates.^28^ Similar variations in the *E*°s were observed in the presence and absence of O_2_ which are characteristic of R and T states.

A combination of quantum chemical calculations and MD simulations have been used to estimate Marcus ET parameters (λ, ΔG°) and ET rates for many redox proteins.^29–34^ These methods make use of the linear response approximation to estimate the free energy parabolas which allows the estimation of λ, and ΔG° from the thermal averages of the energy gaps (<ΔE>) for the oxidized (O) and reduced (R) states.^31^ ET rates can then be predicted using either the full estimation of electronic coupling matrix and Frank-Condon factors or using empirical parameters for protein packing density along with the Marcus parameters.^35,36^ Studies with other redox proteins have shown that mutations both within and outside the active sites lead to significant changes in the Marcus parameters which ultimately modulate the ET rates and protein functions.^35,36,37,38^ MD simulations for the native Hb have been reported earlier.^39–45^ These, mostly focused on the allosteric mechanisms of ligand binding and T → R transitions. More recently Case and Samuel have studied the dynamics of Hb M and associated Heme loss.^46^ But the measurement or estimations of the Marcus parameters and ET rates for Hb and its variants using all atom MD simulations have not been reported till date and represent gaps in our understanding of how globin mutations influence Hb oxidation, reduction and Hb M content via modulation of ET parameters. Reliable predictions of ET parameters for Hb and its variants can provide atomic level and functional insights into factors influencing Hb oxidation. This information can prove valuable in identifying potential Hb variants with increased propensity for oxidation and Hb M formation. It may also prove useful in the design of novel alternative therapeutic interventions in populations where classical therapy (e.g. methylene blue) is contraindicated. Indeed the design of safer blood substitutes has been attempted, but clinical development has often been hampered by Heme-oxidation induced toxicity.^47,48^

In this work, we performed all atom MD simulations on the wild type and variant α Hb chains for the estimation of Marcus ET parameters (λ, ΔG°), and redox potential (*E*°). We address the following key questions with respect to estimation of Hb oxidation among selected Hb variants. 1) What is the influence of Hb M stabilizing α chain mutations on the globin structure and stability? 2) Can a protocol utilizing quantum chemically derived parameters and all atom MD simulations provide insights into the experimentally observed Marcus parameters (λ, ΔG°) for wild type α Hb chain and its redox partner Cyt b5? 3) What is the influence of Hb M stabilizing mutations on the ET parameters? 4) How do these mutations affect outer-sphere (including protein and solvent) reorganization energies (namely, and λ_os_ = λ_prot_ + λ_solv_) and the driving force (ΔG°)?

## 2. Methodology

### 2.1. Protein preparation

Hemoglobin (Hb) α chain coordinates were extracted from T state structure 1HGA (A chain).^49,50^ This structure represents the deoxy-Hb in the T state. As seen in Figure 1 alignment of the protein backbone of this structure with the oxidized methemoglobin (Hb M, 1HGB shows very similar structure (backbone and all atom RMSD = 0.186, 0.233 Å). Thus 1HGA structure was chosen for further analysis. The protonation states for the protein chain were estimated at pH = 7.4 with H++ 3.0^51^ via its webserver.^52^ Total charge on the globin chain was +1 at this pH. For Cyt b5 modeling, the 3NER^53^ structure was selected and protein was prepared for MD simulations using a procedure similar. Total charge on the Cyt b5 structure was -10 at pH = 7.4.

**Figure 1.**
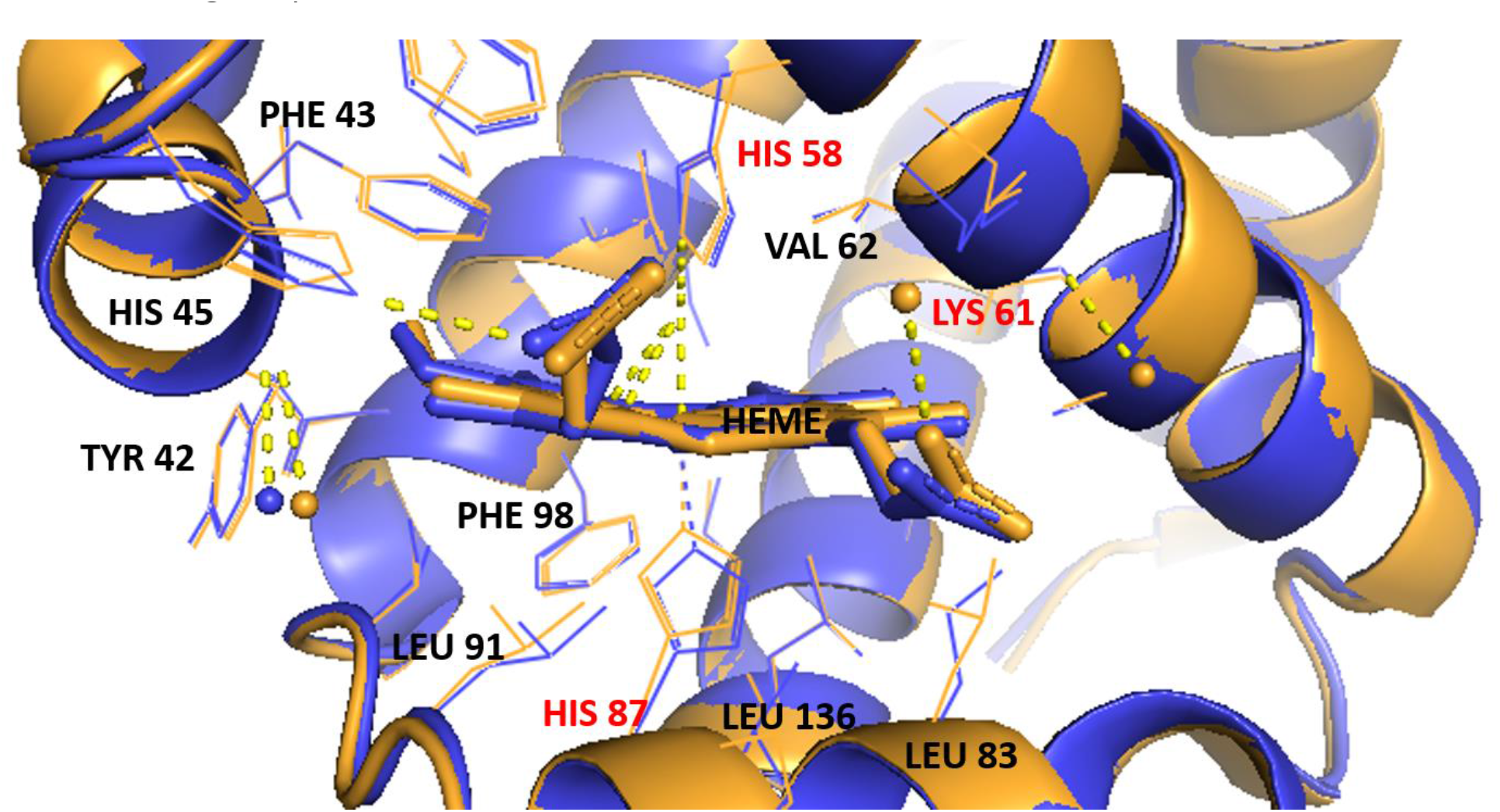
Alignment of α chains of hemoglobin structures in the reduced (1HGA, blue) and oxidized (1HGB, light orange) states. The Fe-N(HIS 87) distances in 1HGA and 1HGB are 2.214 and 2.153 Å respectively. The corresponding Fe-N(HIS 58) distances are 4.444 and 4.303 Å. These parameters are in agreement with the geometry of Heme complexes where the Fe is displaced out of the plane of the porphyrin ring unless coordinated with additional axial ligands like O_2_ and the oxidized state is low spin hexa coordinate usually interacts with water or with HIS 58. Dashed light yellow lines indicate that residues interact with each other. Amino acids in red-color indicate mutation sites studied in this work.

### 2.2. Heme parameterization with MCBP.py and quantum chemical calculations

Metal Center Bond Parameter python tool (MCBP.py)^54^ available in AmberTools18^55^ was used for Heme and axial ligand parameterization in both the redox states of Cyt b5, Hb and Hb variants. A procedure similar to that described for the CYP450 BM3 (Amber advanced tutorial http://ambermd.org/tutorials/advanced/tutorial20/mcpbpy_heme.htm) was used (see Supporting information for details).

### 2.3. Equilibration and production simulations

A robust ten-step protocol recently reported by Roe and Brooks was used to equilibrate the structures of Hb α chains, its variants, Cyt b5 and Cyt b5:Hb complex studied in the present work.^56^ Briefly, step 1 involved minimization of solvent while keeping strong (5.0 kcal/mol Å) restraints on the protein, porphyrin ring, Iron and the axial ligand. In Step 2, a short 15 ps MD with NVT, weak-coupling thermostat was performed while retaining the restraints. Step 3, was 1000 steps of minimization with weaker (2.0 kcal/mol Å) restraints on the same atom selection. Step 4, continued minimization for 1000 steps with even weaker (0.1 kcal/mol Å) restraints on the same atom selection. Step 5, final minimization for 1000 steps without any restraints. Step 6 was MD for 5 ps, with moderate (1.0 kcal/mol Å) restraints on the same atom selection using NPT ensemble. Step 7 and 8 were additional relaxation for 5 and 10 ps respectively with NPT and smaller (0.5 kcal/mol Å) restraints. Step 9, involved unrestrained relaxation for 10 ps. The final density equilibration was performed in 1 ns increments without any restraints under periodic boundary conditions, hydrogens were restrained with SHAKE on, a collision frequency was set to 5 ps^−1^ as recommended for Langevin thermostat and temperature was set to 300 K. This protocol defines three parameter values as a robust criteria for declaring the equilibration success. The criteria are 1) the value for the slope of the density vs. time plot should be less than 10^−6^. 2) The final density of the system should not differ from the average of the second half of the density data by more than 0.02 g cm^-3^. 3) The fitted exponential chi square value should be less than 0.5. All the structure simulated in the present work passed these criteria and thus were used for production runs and further analysis. This protocol available via AmberMDPrep shell script requires the latest version of cpptraj (V4.30.2) which was installed from GitHub and sourced before running the protocol.^57^

The structures for the Hb variants were prepared by renaming the chosen backbone residue names and deleting the sidechain atoms from the PDB file. The mutant side-chain rotamers without clashes and highest probability were selected from the Dunbrack’s rotamer library available in Chimera.^58,59^ Identical protocols were used for equilibrating the Hb variant α chains in both redox (Fe^2+^ and Fe^3+^) states. The summary of mutation categories in the Hbvar database contains information on 363 α chain variants. The summary table also gives 13 variants classified as Methemoglobins, 4 of which are α chain variants. Nonetheless, many of the 363 α chain variants (not classified in the database as Methemoglobins), do form Hb M. Thus we additionally selected variants which have been characterized in the literature^60^ and are known to form methemoglobin (Hb M) and/or lower the concentration of reduced Hb namely, Hb M Boston (H58Y), Hb M Iwate (H87Y), Hb Miyagi (K61E) and Hb Kirklareli (H58L). Additionally, a single point mutant which does not lead to Hb M formation, Hb J-Buda (K61N), was studied.

## 3. Results and Discussion

Reduction of the monomeric native α Hb chain has been studied earlier.^62,63^ These studies found that the redox behavior of the α Hb chain in monomeric state is almost identical to the Hb tetramer. Thus studying the effect of α Hb monomer on the Marcus parameters is justified and can be expected to meaningfully represent the ET and redox processes in the tetramer. As noted by Hub et al., most of the experimental studies monitor the hydrogen bond dynamics between a small set of selected inter-chain residues e.g. αAsp94-βTrp37 and αTyr42-βAsp99.^41^ These interactions are interpreted in terms of conformal (T ↔ R) transitions without further investigations. NMR studies in solution have identified α chain His45, Tyr42, Thr41, His89, His50, His72 residues play an important role in Bohr effect while residues around Asp94 and His122 participate in α1β2 subunit interactions.^17^

With this background the results and discussion is divided into four main subsections; 3.1. Influence of Hb α-chain mutations on the structure and stability, 3.2. Marcus ET parameters and redox potential (E°) for the wild type Hb, 3.3. Marcus ET parameters (λ, ΔG°) and redox potential (*E*°) for selected Hb variants, 3.4. Contributions of protein and solvent to the reorganization energies, 3.5. Marcus ET parameters for the Cyt b5 (the redox partner of Hb).

### 3.1. Influence of Hb α-chain mutations on the structure and stability

Realizing the dynamical nature of Hb α chain interactions the effect of Hb M stabilizing mutations on the dynamics and flexibility of various globin regions was investigated. Residue wise backbone root mean square fluctuations (RMSF) on the equilibrated trajectories (40 ns) were calculated with cpptraj (as seen in Figure 2). As expected the N and C-terminals show higher fluctuations. A comparison of the RMSF for residues ∼10-130 shows that the overall globin structure is similar among variants except the Hb M Iwate which involves mutation of the axial Histidine ligand with Tyrosine (H87Y). This mutation is known to destabilize the Heme globin structure and function.^64^

**Figure 2.**
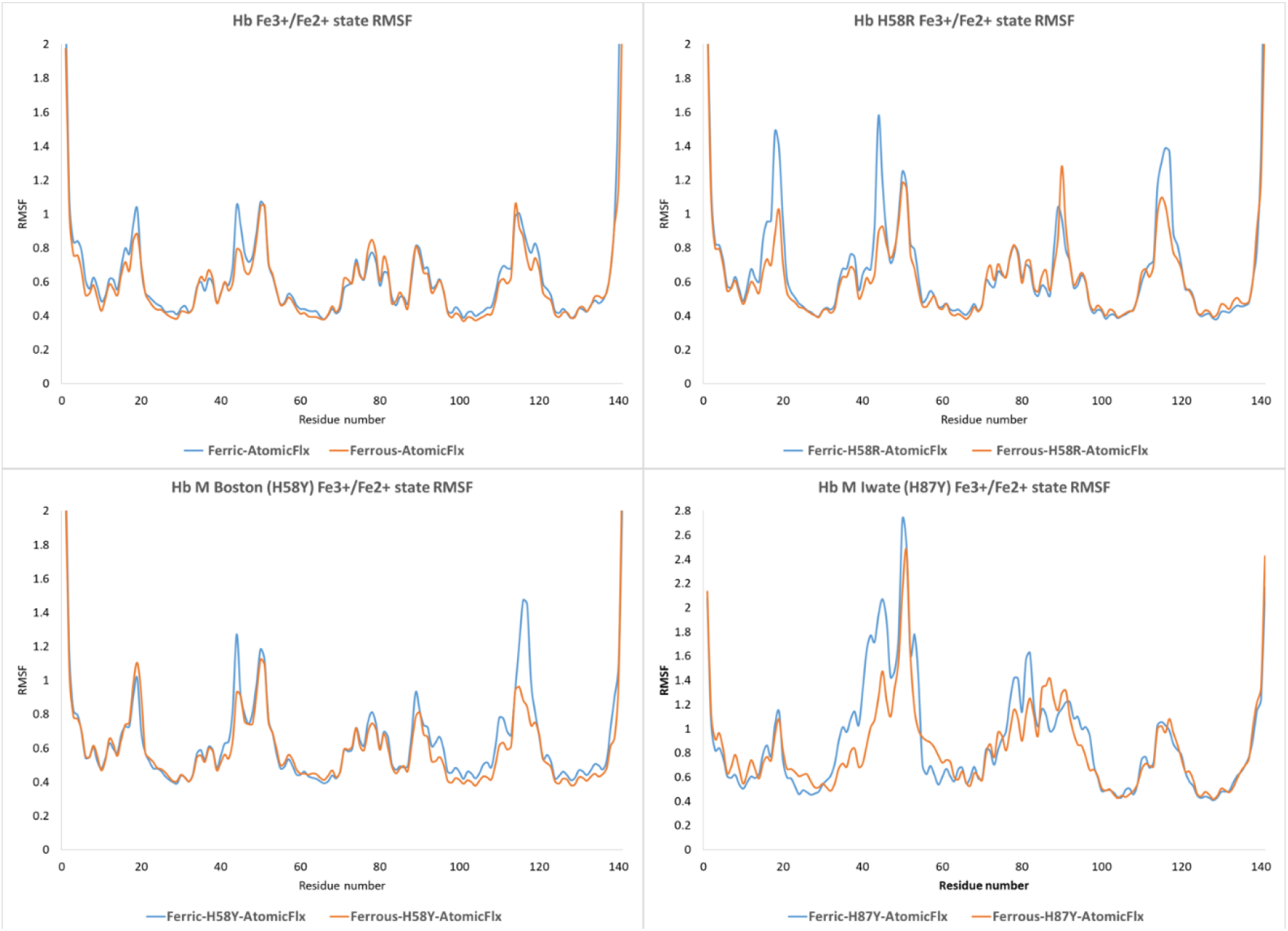
RMSF for the protein backbone for the wild type Hb, mutants leading to the formation of Hb M (Iwate and Boston variants) and H58R mutant which is analogous to the β chain Hb Zurich variant. The RMSF was calculated for 2000 snapshots extracted from a 40 ns MD trajectory.

To assess the influence of other mutations which show a stable globin structure but nonetheless oxidize to Hb M, the RMSF values among their redox states were compared. For this a student T-test matrix among all the variants was calculated (See Table S1, Supporting Information). Residue-wise RMS fluctuations (RMSF) are considered similar for pairs that show a P value more than 0.05. Mutation of the distal Histidine with Tyrosine (H58Y) seen with the Boston Hb variant maintains the overall globin structure. The largest backbone fluctuations are in the residues 44-54, 89, 90, and 114-119 (see Figure 2). The region 44-54 is a loop that has important hydrophobic (Phe43, 46-Heme) and electrostatic (His45-Heme carboxylate) interactions that stabilize the Heme in the wild type. Whereas the region 114-119 has key interactions with the β chain residues (Arg30, and His116). The RMSF (excluding four terminal residues) for the redox state between native and H58Y variant shows that fluctuations are significantly different and the average, min, max and stdev are larger for the H58Y variant (Table S1 and S2, see supporting information). This indicates that this mutation destabilizes both the redox states.

Mutation of the His58 to Arg, analogous to the H58R Zurich variant of the β chain, shows similarly large variations in these regions in addition to a larger fluctuation in the loop 117-120. These findings are consistent with the large size of the Arg side chain (similar to Tyr) which leads to similar motions of these regions as manifested by higher RMSF value (see Figure 2). The fluctuations induced by H58R mutation are significantly different and average values are about twice from the wild type Hb (Table S1 and S2).

A close inspection of the globin and Heme structure in Hb M Iwate during the MD simulation, shows that the Heme group significantly moves out of the protein pocked and gets exposed to the solvent medium (see Figure S1). It is known in the literature that the Hb M Iwate structure is not stable and there is experimental evidence that the Heme group gets transferred to the distal histidine in the reduced state.^64^ Considering the large structural changes that H87Y mutation induces in the Hb structure and the lack of crystal structure we choose not to use these MD trajectories to investigate the effect on Marcus parameters, oxidation activation energies and ET rates for this variant.

The RMSF analysis for the Hb Miyagi and Kirklareli variants (K61E and H58L) which are known to form Hb M showed similar RMSF (pairwise) compared to the reduced (Fe^2+^) and oxidized (Fe^3+^) native Hb. The average reduced (Fe^2+^) RMSF is larger than the oxidized (Fe^3+^) state for both the variants (Table S1 and S2). This is in contrast to the native and other Hb variants suggesting that the reduced state fluctuations are higher in these variants. Recently a crystal structure (3QJD) for the Kirklareli variant has been reported.^65^ Pairwise CA atom RMSD between the 1HGA and 3QJD is 1.00 Å, suggesting that the mutant adopts a similar globin structure (see Figure S3). MD simulations starting from this structure showed similar trends in RMSF. Nonetheless, the Fe^2+^ state showed larger fluctuations for polar residues Asn78, His89 and Lys90. This suggests that the His58 interactions with Heme stabilize the movement of proximal residues and mutation to a non-polar Leucine thus increases the dynamics of these residues. The Fe^3+^ state, on the other hand, had higher RMSF in the loop region (residues 114-117). In contrast the Hb J-Buda variant with a semi-conservative mutation (K61N) which showed minor fluctuations in the Heme interacting loop (residues 44-54). Whereas relatively larger differences in fluctuations are observed in the region 115-120 which indicates weaker interactions with the β chain (especially in the oxidized state, see Figure 3). Except for the K61E, H58L variants, the RMSF values are consistently higher for the oxidized (Fe^3+^) state. This can be understood by considering the reduction in the electrostatic repulsion between the Glu side-chain and the Heme group in the oxidized state. These findings suggest that mutation in one region of the Hb can have significant influence on the dynamics in distant regions.^66,67^

**Figure 3.**
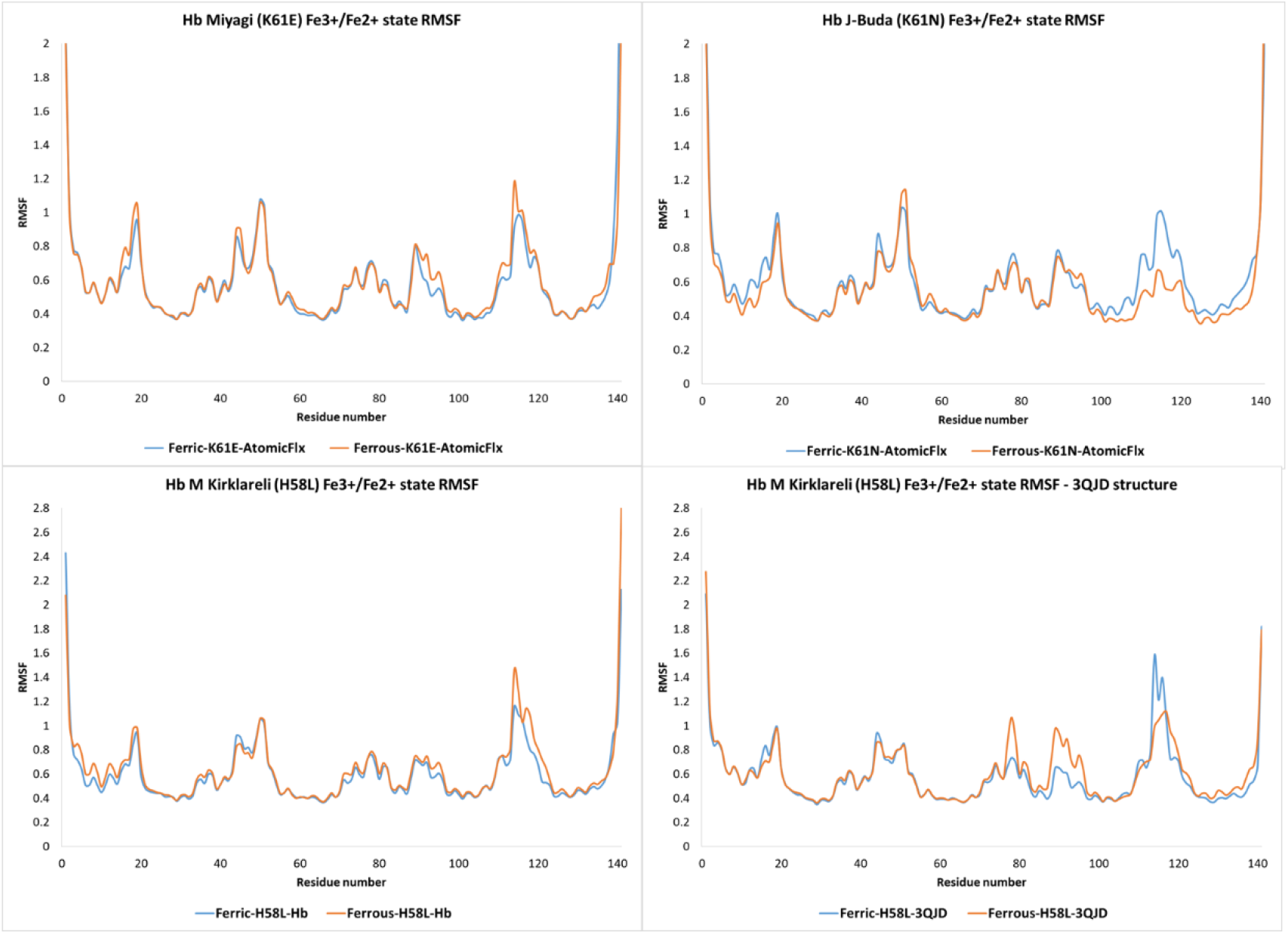
Residue wise RMSF for the protein backbone for the Hb variants in the oxidized (Ferric) and reduced (Ferrous) states during the 40 ns MD simulations. The Hb Miyagi (K61E) and Hb Kirklareli variants forms Hb M, while the Hb J-Buda (K61N) variant functions normally.

A comparison of the RMSD for the Hb redox states similarly shows that the variant leading to Hb M formation results in the largest movement in the protein backbone (see supporting information Figures S1-S9).

### 3.2. Marcus ET parameters and redox potential (*E*°) for the wild type Hb

Since different mutations lead to different Hb reduction rates and different equilibrium Hb M content, we tested if there is a correspondence between these and the Marcus ET parameters. As mentioned in the introduction MD trajectories have been used earlier along with the linear response approximation to determine Marcus ET parameters (λ, ΔG°) and ET barriers for redox proteins. Similar to such proteins, in the Heme proteins small changes in system charges are distributed over large number of atoms (porphyrin ring, axial His, and residues within non-bonded interaction distances). For example the inner sphere contribution to reorganization energy was estimated to be only 50 meV in cytochrome MtrF (a decaheme bacterial redox protein complex).^68^ Similar to Hb, the Cyt MtrF Hemes are partially solvent exposed and are known to exhibit λ in the range 0.75–1.1 eV. Thus it is reasonable to expect that the variations in the structure and dynamics of the Hb Hemes will contribute only a small fraction (< 5 %) to the total reorganization energy (i.e. λ >> λ_is_, where subscript “is” represents inner coordination sphere of the metal center). The most important contributions come from the protein (λ_prot_) and the solvent (λ_solv_) environments which together constitute the outer sphere reorganization energies (λ ∼ λ_os_ = λ_prot_ + λ_solv_). Thus these λ values were estimated by taking the thermal averages of the AMBER force field energies calculated by the post-processing of MD trajectories with cpptraj. The sander program (parallel version) was used to calculate the energy gaps in periodic conditions similar to those used for MD simulation. The λ_prot_ was obtained by stripping water molecules and counter ions from the trajectory and recalculating the AMBER force field energies.

The mutations studied in this work are within the Hb active site and previous docking models of the Hb-Cyt b5 complexes have estimated a Heme edge to edge distance of ∼ 8 Å.^69^ Thus it is reasonable to expect that the relative Hb M reduction rates and content would correspond to relative changes in Hb redox properties. Since Hb oxidation to Hb M is known to occur via multiple enzymatic or non-enzymatic steps, the redox process can be simulated as a one-step oxidation process for a single ionizable group shown in Equation 2.^32^ Using this formalism and the linear response approximation the λs are equated to the half of the differences in the average vertical ionization energies (<Δ*E*_*M*_>, called the average energy gaps) for the redox states (M = R, O). These average energy gaps are calculated using <ΔE_M_> = <E_O_> - <E_R_>, where <E_M_> are the Boltzmann average potential energies of the M state calculated using AMBER force field parameters for the O and R redox states respectively.^32^ The free energy changes (ΔA° ∼ ΔG°) are equated to the mean of these energies gaps (see equations 2-19 in reference ^32^ and equations 13-15 in reference ^31^).

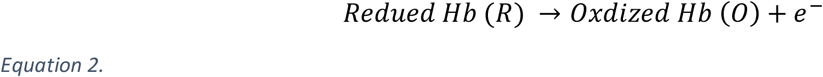

Table 1 below shows the average vertical ionization energies (<ΔE>) for the redox states, Marcus parameters (λ, ΔG°), and the redox potential (*E*°). Since the non-polarizable force fields are known to systematically overestimate reorganization energies, a factor of 1.6, recommended by Blumberger was used to scale λ values.^31^ The scaled λ are discussed here onwards (except in section 3.4, where relative changes in protein and solvent contribution are discussed).

**Table 1.** The total reorganization free energies (λ in eV, scaling factor 1.6),^31^ redox potential (E°, V), for Hb, Cyt b5, and selected α chain Hb mutants.

| Protein | Mutation | $\lambda$ | $E^\circ$ | Hb M formation |
| --- | --- | --- | --- | --- |
| Hb | Wild | 0.6853 | -0.165<sup>a</sup> | < 1 % |
| $\alpha$ analogue of the Hb Zurich $\beta$ variant | H58R | 0.7216 | -0.064 | NA |
| Hb Miyagi | K61E | 0.7297 | -0.296 | Increased Hb M formation |
| Hb J-Buda | K61N | 0.6692 | -0.204 | NA |
| Hb Kirklareli | H58L | 0.7001 | -0.176 | Increased Hb M formation |
| Hb Kirklareli (3QJD) | H58L | 0.6867 | -0.150 |  |
| Cyt b5 (3NER) | Wild | 0.9299 | -0.212 | Not applicable |
| <sup>a</sup> Computed redox potential ($E^\circ$) shifted to match experimental $E^\circ$ versus normal hydrogen electrode. Same shift constant used for all mutants. | | | | |
| <sup>NA</sup> Information not available in the Hbvar database and literature. |  |  |  |  |

The wild type Hb showed vertical ionization energies (ΔE) of 0.91 and 3.10 eV in the oxidized and reduced states respectively (see Table S3). This gives a reorganization energy (λ) for the wild type Hb 0.6853 eV, which is lower than the value reported by Blankman et al., for the Hb tetramer.^28^ It should be noted that the values for the Hb tetramer include λ contributions from the β subunit and is known to undergo larger structural changes upon ligand binding and hence increase the λ in the tetrameric complex (Figure 8a, 10 of reference ^40^).

Moreover our estimates for α Hb, λ are slightly lower than Heme complexes with relatively higher solvent exposure.^68^ These agreements in the calculated ET parameters with the experimental data lend support to our modeling approach.

As mentioned above a redox potential (*E°* vs NHE) of -0.165 V has been reported in the literature for the T-state Hb.^27^ Thus ΔG° predictions for Hb were scaled to match the experimental data. Further using these values a good estimate for the biologically relevant ET from Cyt b5 to Hb can be modelled using the computed Cyt b5 λ = 0.9299 eV (see section 3.5). The experimental electrochemical λ = 0.44 eV^28^ is smaller than reported for similar proteins earlier^32^ most probably due to adsorption of the Cyt b5 on the electrode surface which reduces the protein motion significantly. Thus the relative changes in the Marcus ET parameters and redox potential in different Hb variants with and without an increased tendency to form methemoglobin (Hb M) are discussed in the next section.

### 3.3. Marcus ET parameters (λ, ΔG°) and redox potential (*E*°) for selected Hb variants

The variants resulting from the mutation of residues in the ligand binding sites are known to affect O_2_ affinity, Hb function and often increase the Hb M content.^2^ Thus we investigated the influence of selected mutations on ET parameters, oxidation free energies and redox potential (*E*°). The mutation of the distal Histidine with Tyrosine (H58Y) leads to the Hb M Boston variant. It shows larger λ (∼ 0.792 eV) but a similar free energy change (ΔG°) and *E*° compared to the wild type which might make its experimental assignment difficult. Raman spectroscopic studies on the Hb Boston and Iwate variants have shown that these variants involve the coordination of the Heme with distal ligands which was not modelled explicitly here.

Nonetheless, modern force field based MD simulations are expected to capture the out sphere effects on ET parameters. The influence of the mutation of distal Histidine to a non-coordinating Arginine residue (H58R) on ET parameters and redox potential was studied. As seen in Table 1, this variant shows larger λ (0.7216 eV), and ΔG° (2.105 eV) values giving higher redox potential (*E*° = 0.064 V) than the wild type Hb. These parameter values indicates a lower equilibrium concentration of Hb M for this variant. Although, this variant has not been reported in the literature yet, its β analogue, the Hb Zurich variant, has been found to be susceptible to oxidation (HbVar database). Our results suggest that this Hb (H58R) variant may not lead to a noticeable change in the background Hb M content. In contrast the Hb Kirklareli variant has been reported to undergo oxidation to Hb M (HbVar database).^65^ This mutant involves the substitution of His58 with a Leucine. Our calculations showed an increase in the reorganization energy (λ = 0.7001 eV), and free energy (ΔG = 1.993 eV) of oxidation similar to the wild type Hb. The corresponding change in the estimated *E*° is < 10 mV. MD simulations with the recently reported crystal structure for this variant (3QJD),^65^ gave slightly different ET parameters (λ, ΔG°). The calculations using 3QJD structure give a slightly higher energy gap (ΔE) for the oxidized state (Table S3), and while it remains similar for the reduced state. The λ decreases by 0.0134 eV, ΔG° increases to 2.02 eV, thus giving a lower *E*° = -0.150 V, suggesting higher equilibrium concentration of oxidized Hb (i.e. Hb M) compared to the wild type Hb. Thus for the Hb Kirklareli variant using the crystal 3QJD structure seems necessary to make accurate predictions of relative ET parameters that agree with the experimental information.

Another active site residue (LYS61) forms key H-bonding interactions with the Heme carboxylates and contributes in stabilization of the reduced (Fe^2+^) state. Mutation at this position into a GLU residue forms the Hb Miyagi variant. This variant (Table 1 and Table S3) shows relatively larger ΔE and λ values, while a lower ΔG° and *E*°= 1.87 eV and -0.296 V are predicted respectively for the Hb oxidation. This suggests a higher equilibrium concentration of Hb M content in agreement with the experimental fact (HbVar database). On the contrary the Hb J-Buda variant involves a mutation of this LYS61 to a semi-conservative ASN residue and exhibits a lower λ = 0.6692 eV, and the ΔG° = 1.97 eV is closer to the wild type, while the predicted redox potential (*E*°) = -0.204 V is lower than wild type and higher than the Hb Miyagi variant.

### 3.4. Contributions of protein and solvent to the reorganization energies

As mentioned earlier, the total reorganization energy (λ) comprises of the inner-sphere and outer-sphere contributions. Similar to cytochromes, the relatively rigid nature of the Heme cofactor, causes the inner-sphere reorganization energies to be small for Hb and variants with mutations in the outer-sphere protein environment.^70^ Table 2 shows the protein reorganization energies (λ_prot_), calculated by stripping the solvent and ions from MD trajectories and re-estimating the AMBER energies with ff19SB force filed. The solvent reorganization energies (λ_solv_) are then estimated as the difference between the total and protein reorganization energies (i.e. λ_solv_ = (λ - λ_prot_)). Since the relative differences in these energies are more relevant we discuss the unscaled λ values. For all the Hb variants including the wild type, the major contribution to total λ comes from the solvent reorganization (λ_solv_: 0.514 to 0.706 eV, 73-97 %). For the wild type Hb, the solvent contributes 76 %, which remains constant for the Hb M Boston variant. Nonetheless, the λ_prot_ are non-negligible for the wild type (0.162 eV, 24 %) and the Hb M Boston variant (0.179 eV, 25 %). The H58L (Hb Kirklareli) variant showed a larger protein reorganization (λ_prot_ = 0.186 eV, 27 %) and solvent reorganization (λ_solv_ = 0.514 eV, 73 %) comparable to the wild type Hb in response to oxidation.

**Table 2.** The average vertical ionization energies (<ΔE>) for the protein redox states (solvent molecules were stripped before single point energy estimation with sander), the protein and total reorganization free energies (λ_prot_, and λ), solvent reorganization energy (λ_solv_) for Hb wild and selected mutants. The relative differences in λ, and its protein, solvent contributions between the Hb wild type and variants are important.

| Hb $\alpha$ chain Variants | Mutation | $\lambda$ | $\lambda_{\text{prot}}$ | $\lambda_{\text{solv}} = (\lambda - \lambda_{\text{prot}})$ | % contribution of $\lambda_{\text{solv}}$ to $\lambda$ |
| --- | --- | --- | --- | --- | --- |
| Hb | Wild | 0.685 | 0.162 | 0.524 | 76.43 |
| Hb M Boston | H58Y | 0.693 | 0.173 | 0.520 | 75.02 |
| $\alpha$ analogue of the Hb Zurich $\beta$ variant | H58R | 0.722 | 0.179 | 0.543 | 75.21 |
| Hb Miyagi | K61E | 0.730 | 0.023 | 0.706 | 96.81 |
| Hb J-Buda | K61N | 0.669 | 0.032 | 0.638 | 95.28 |
| Hb Kirklareli | H58L | 0.700 | 0.186 | 0.514 | 73.38 |

The H58R variant shows λ_solv_, and λ_prot_ values similar to the wild type. In the wild type Hb, the sidechain of LYS61 forms an average of two H-bonding interactions with the bulk water and one H-bond with Heme carboxylates (see Figure 4). These H-bond interactions between residue 61 and Heme are lost in the K61E Hb Miyagi variant. The glutamate residue in the K61E Hb variant now forms an average of 5.8 and 4.3 h-bonds with the bulk water molecules in the reduced (Fe^2+^) and oxidized (Fe^3+^) states respectively. The repulsion between the glutamate and Heme carboxylate leads to a significant reorganization of solvent structure (λ_solv_ = 0.706 eV, 97 %) forced by increased H-bonding with the surrounding water molecules. The Lys61 residue of the wild type Hb forms H-bonds with the Heme in 26 % of the frames and thus stabilizes the additional negative charge in the reduced (Fe^2+^) state (0.07 % of the frames in oxidized, Fe^3+^ state). This reduces to negligible (0.0075 % of the frames) in the K61N variant and zero in the K61E variant. This explains the unexpectedly lower protein reorganization in these variants in response to oxidation. This suggests that the Lys61 residue place a key role in determining the protein reorganization in response to oxidation and mutation at this site (especially into a negatively charged amino acid) increases Hb M formation. The larger protein reorganization (λ_prot_) in the wild type prevents faster oxidation to Hb M, but mutation of Lys to uncharged or negatively charged residues lowers the λ_prot_ facilitating Hb M formation. Although there is a corresponding increase in the λ_solv_, this is likely to be spread out among the a large number of bulk water molecules thus keeping the activation free energy lower than the wild type Hb.

**Figure 4.**
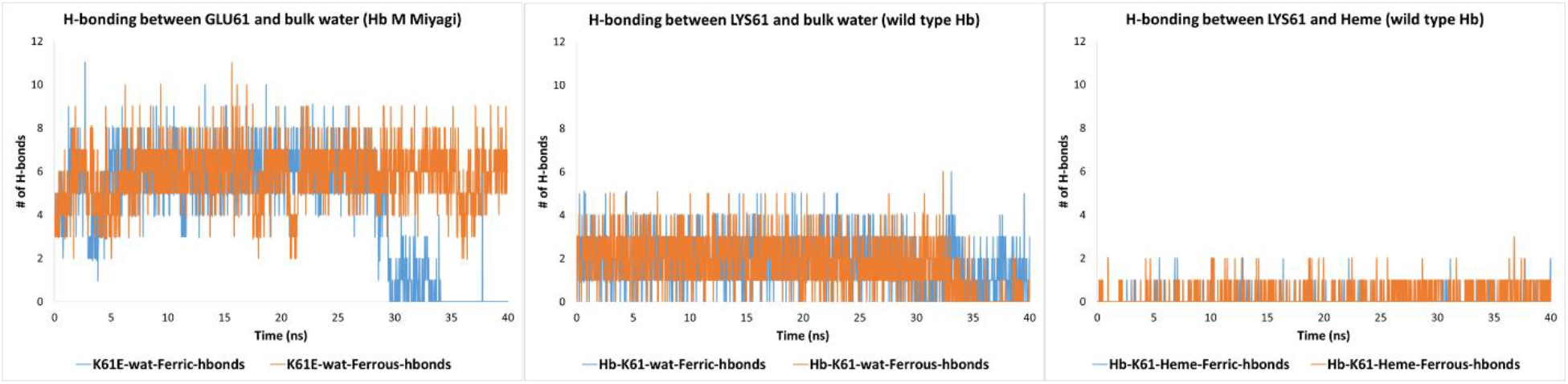
Number of H-bonds formed between the side chain of the residue 61, solvent water and Heme carboxylate in the wild type Hb (K61) and in the Hb M Miyagi variant (K61E) during the 40 ns MD simulation.

The Hb J-Buda variant which involves a semi-conservative mutation (K61N) forms an average only 1.1 H-bond interactions with the bulk water (not shown), and no interactions with the Heme group. This explains lowest average RMSF calculated for both the K61E and K61N variants (Table S2). Nonetheless, there is a difference of 0.068 eV in λ_solv_ between these two variants with the K61N (Hb J-Buda) being similar to the wild type. This coupled with the larger oxidation free energy difference (0.13 eV) for the Hb Miyagi (K61E) variant also offers logical explanation for its lower redox potential.

### 3.5. Marcus ET parameters for the Cyt b5 (the redox partner of Hb)

As mentioned in the introduction Cyt b5 reduces the oxidized Hb (Hb M) under physiological conditions i.e. within RBCs. Thus it is important to understand the ET properties of this important redox partner of Hb. Only one Crystal structure for only one human Cyt b5 has been reported (3NER).^53^ Thus this structure was used to model the ET properties of the Hb redox partner. Protein preparation, Heme parameterization and MD simulations were performed using the protocol mention in section 2. Table 1 shows the ET parameters (λ, ΔG°) for the oxidation of Cyt b5. The energy gaps for the Cyt b5 are considerably larger than those predicted for Hb and its variants (see Table S4). This is probably due to the smaller size and associated larger conformational change in the protein structure upon oxidation. Thus the λ value for Cyt b5 are also larger than Hb and giving a scaled λ = 0.9299 eV. This estimate is larger than the electrochemically determined λ^28^ and half of MD based predictions for Cyt b5/Cyt c complex reported earlier.^71^ This is expected since the electrochemical adsorption and protein complexation significantly stabilizes the protein dynamics in Cyt b5 in comparison to isolated solution state. The calculated ΔG° is also higher and was scaled to match the experimental redox potential of -0.212 V.

## Conclusions

In summary, the answer to the questions asked in the introduction are as follows. 1) The single point mutations studied here, mostly do not disrupt the globin structure. Major exceptions to this are the His → Tyr mutations (Hb M Boston; distal site and Hb M Iwate, proximal). In the later variant a Heme transfer to Tyr is reported in the literature. Mutants K61E, K61N, H58R and H58L show influence on the selected regions of the globin chain (loop 44-54 and region 114-120). 2) All atom MD simulations and average vertical energy gap calculations allow reasonably accurate prediction of Marcus parameters (λ, ΔG°) that are in agreement with literature reports on similar Heme proteins. 3) All the Hb M forming mutations (H58Y, K61E and H58L) lead to an increase in the total reorganization energy (λ) associated with the Hb oxidation. Whereas the Hb J-Buda (K61N) shows a marginal decrease in λ. The calculated redox potentials are in correspondence with the known propensity of these variants to form Hb M. 4) The solvent reorganization (λ_solv_) makes the largest contributions for all the variants (including the wild type). The mutation of the Lys residue has the largest influence on the protein reorganization energies (λ_prot_) thus highlighting its role in the stabilization of the reduced (Fe^2+^) state, normal Hb structure and function. 5) The calculated Marcus ET parameters, redox potentials offer explanation for the corresponding higher and moderate propensity of Hb oxidation to Hb M for the K61E and K61N mutants. Our calculations are also consistent with the reports that H58L variant (Hb Kirklareli) undergoes faster autoxidation (lower *E*° than wild type Hb).

Finally, this study represents the first attempt to calculate Marcus parameters for the oxidation of Hb to Hb M using all atom MD simulations of the redox states. Our GPU enabled MD simulations, post-processing, calculations and data analysis completes within 4-5 hour/mutant. Thus this methodology can be applied to other Hb variants and Heme proteins to extract meaningful predictions and has potential to guide experimental studies on most interesting and useful mutations.

## Author contributions

VAD conceptualized the project, wrote funding proposal, performed literature search, calculations, analyzed results, wrote the manuscript and participated in discussions. JB analyzed the results, provided expert opinion, participated in manuscript writing and discussions. SKV performed preliminary literature and database search, and participated in discussions.

