## supporting information for "Electron transfer parameters for Methemoglobin formation in mutant Hemoglobin α-chains"

Figure 1. Heme-axial HIS87 models used for defining atomic charges and metal center bonded parameters. A) Small model of the metal center used for optimization and frequency calculations. Atoms marked with an asterisk Symbol (*) were frozen during optimization to avoid formation of artefactual H-bonding interaction between the NH proton and carbonyl oxygens. B) Large model of the metal center used for defining chargers and atomic radii. ........................................................................ 3

Figure S2. Hb structures (reduced state) extracted from the MD trajectories for the two Histidine to Tyrosine variants namely Hb M Iwate (H87Y) and Hb M Boston (H58Y). The structures after initial minimization (red), equilibration (green), at 20 ns (blue) and 40 ns (magenta) are shown for both structures for comparison. RMSD for the Hb M Iwate structures are > 1.2 Å, while for the Hb M Boston structures it remains below 1.0 Å (w.r.t. to the minimized structure). The sphere represent the Iron atoms. A similar pattern was observed for the oxidized state structure (not shown here)....................... 4

Figure S3. Active site view of the backbone aligned 3QJD and mutated 1HGA (H58L) structures. The 1HGA structure mutated at 58 residue has been equilibrated after selecting the most likely rotamer selected from the Dunbrack's library. The CA RMSD between these two structures is 1.0 Å while the all atom RMSD is 1.1 Å. This shows significant differences in the orientation of the side chains of the active site residues while the backbones alignment are comparatively similar. LEU58 has been labelled.......... 5

### Heme parameterization with MCBP.py and associated quantum chemical calculations

In brief, 1) Heme group was reduced with reduce 3.3.[1] 2) Atoms were renumbered using pdb4amber. 3) BCC method as implemented in antechamber was used to calculate atomic charges in mol2 format.[2,3] 4) Bonded and non-bonded parameters were extracted using parmchk2. 5) metalpdb2mol2.py script was used to generate FE.mol2 files with +2 and +3 charge for the reduced and oxidized states respectively. The protein structure generated by H++ was merged with porphyrin and Iron coordinates and renumbered for use with MCBP.py tool.

In the input file for MCBP.py tool a cut off of 2.8 Å was used for identifying amino acid residues directly interacting with the Iron. Metal atom number in the protein structure, mol2 files for Iron, porphyrin ring and frcmod file for the latter were also identified in the input file. This was followed by four steps as recommended in the Amber18[4] manual and MCPB.py tutorial. In the first step input files for quantum chemical calculations on the small (Heme + side chain of axial HIS87) and large models (Heme + axial HIS87) were generated (Figure 1). Considering the stability of the high spin states for the unligated Hb,[5–7] the charge/multiplicity for the reduced (Fe2+) and oxidized (Fe3+) states were set to -2/5 and -1/6 respectively. Geometry optimization and frequency calculations were performed on the small model structure using Gaussian 09 program[8] at B3LYP/6-31G(d) level while freezing the HIS87 side chain NH and Heme carbonyl carbons. Charges and atomic radii were estimated using single point energy calculations performed at the same level of theory for the large model.

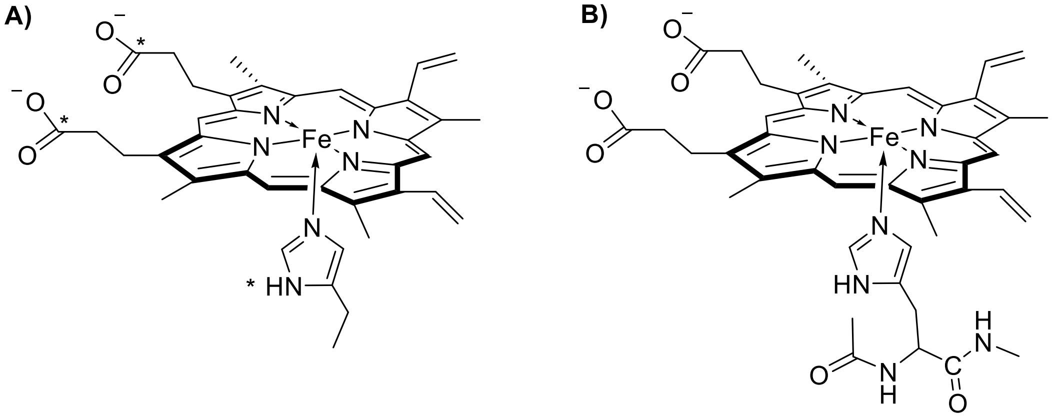

*Figure 1. Heme-axial HIS87 models used for defining atomic charges and metal center bonded parameters. A) Small model of the metal center used for optimization and frequency calculations. Atoms marked with an asterisk Symbol (*) were frozen during optimization to avoid formation of artefactual H-bonding interaction between the NH proton and carbonyl oxygens. B) Large model of the metal center used for defining chargers and atomic radii.*

In the second step with MCBP.py script Seminario method was used to generate force field parameters using the Gaussian 09 output, and formatted checkpoint files. The third step involved RESP charge fitting and mol2 file generation. Partial atomic charges estimated for the Iron atom were 0.475 and 0.607 in the corresponding reduced and oxidized states. The fourth step involved generation of input file for leap which generates the final set of Amber topology, parameters and input coordinate files required for performing MD simulations. This leap input file was edited to load the ff19SB, gaff force field parameters, while Li/Merz monovalent ion parameters for TIP3P water model were loaded.[9,10] A TIP3P water box of 10 Å was added around the protein, along number of counter ions ($Na^{+}$, or $Cl^{-}$) such that the protein with $Fe^{2+}$ redox state is neutral and thus the $Fe^{3+}$ state has +1 charge. With this protocol the prmtop and inpcrd files remain compatible for the estimation of thermal averages of the energy gaps (<ΔE>) using cpptraj and sander programs. The number and type of ions added for each $Fe^{2+}$ system are given in the main text.

The bis-His Heme environment around the Fe in Cyt b5 leads to stabilization of low-spin states, thus hte charge/multiplicity for the reduced (Fe2+) and oxidized (Fe3+) states were set to -3/2 and -2/1 respectively. Rest of the parameterization procedure was similar to that used for Hb.

### Summary of the structural changes in Hb M Boston and Hb M Iwate during MD simulations

A close inspection of the globin and Heme structure in Hb M Iwate during the MD simulation, shows that the Heme group significantly moves out of the protein pocked and gets exposed to the solvent medium. A comparison of the Hb structures for the two Histidine → Tyrosine mutants in shown in Figure S2. The Heme structure for the Hb M Boston variant with mutation of the distal histidine (H58Y) doesn't change significantly after equilibration and during the 40 ns MD simulation. On the contrary, the Heme (and globin) structure for the Hb M Iwate variant has already started to move out of the protein active site during the equilibration phase. The Heme group continues to move out of the protein pocket, gets fully exposed to the solvent and the porphyrin ring becomes orthogonal with the initial plane (red vs magenta structures in Figure S2).

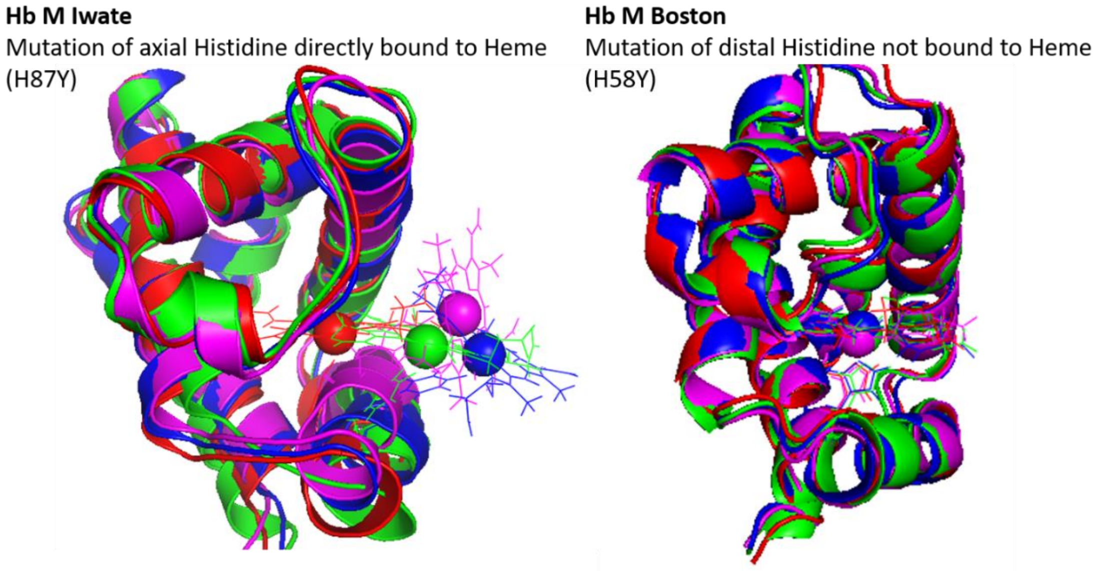

*Figure S2. Hb structures (reduced state) extracted from the MD trajectories for the two Histidine to Tyrosine variants namely Hb M Iwate (H87Y) and Hb M Boston (H58Y). The structures after initial minimization (red), equilibration (green), at 20 ns (blue) and 40 ns (magenta) are shown for both structures for comparison. RMSD for the Hb M Iwate structures are > 1.2 Å, while for the Hb*

*M Boston structures it remains below 1.0 Å (w.r.t. to the minimized structure). The sphere represent the Iron atoms. A similar pattern was observed for the oxidized state structure (not shown here).*

It is known in the literature that the Hb M Iwate structure is not stable and there is experimental evidence that the Heme group gets transferred to the distal histidine in the reduced state.[11] Thus these MD trajectories and structures for the Hb M Iwate probably represent artifacts mostly due to not explicitly considering the electronic structure of the Heme group. Nonetheless, the simulation is consistent with the instability of this variant. Considering the large structure changes H87Y mutations introduces in the Hb structure and the lack of crystal structure we choose not to use these MD trajectories to investigate the effect on Marcus parameters and ET activation energies for this variant.

### Comparison of 3QJD and 1HGA-H58L-rotamer equilibrated structure

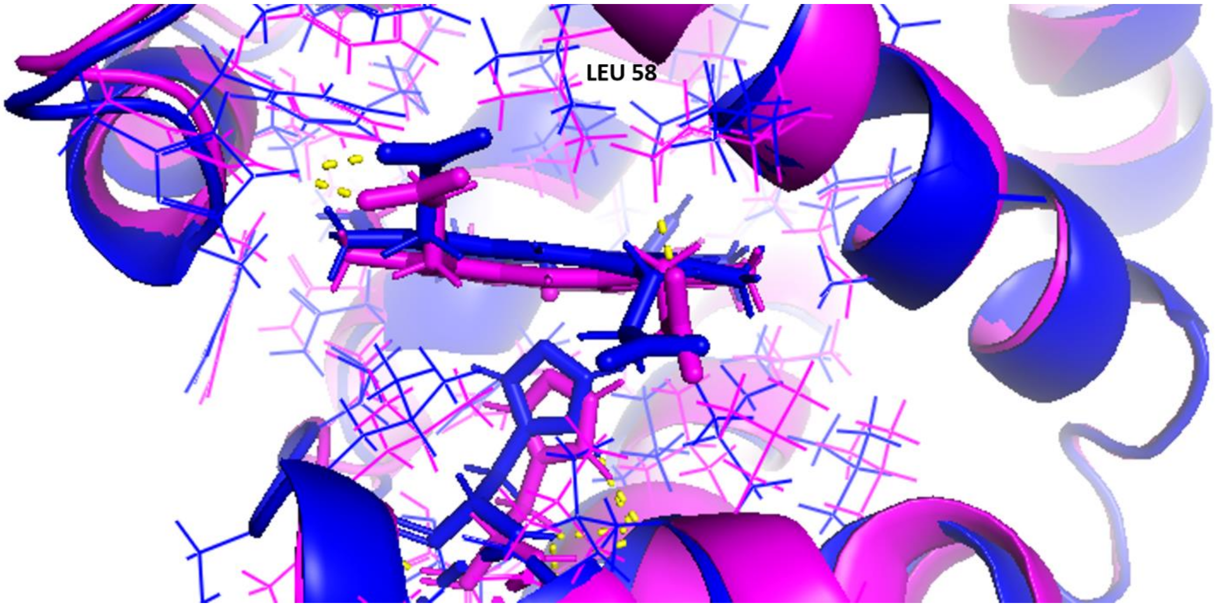

*Figure S3. Active site view of the backbone aligned 3QJD and mutated 1HGA (H58L) structures. The 1HGA structure mutated at 58 residue has been equilibrated after selecting the most likely rotamer selected from the Dunbrack's library. The CA RMSD between these two structures is 1.0 Å while the all atom RMSD is 1.1 Å. This shows significant differences in the orientation of the side chains of the active site residues while the backbones alignment are comparatively similar. LEU58 has been labelled.*

*Table S1. Student T-test P value matrix for the residue-wise RMSF values in the redox (Ferric: Fe3+ and Ferrous: Fe2+) states of the different Hb variants studied in this work. Highlighted cells show pairs of redox states for which backbone RMSF values where similar. A P > 0.05 indicates that the two groups have similar residue-wise RMSF values. Four residues from each terminal have been excluded in this analysis since they are known to generally show larger RMSF value.*

| N/C-ter excluded | Fe3+-1HGA | Fe2+-1HGA | Fe3+-H58R-1HGA | Fe2+-H58R-1HGA | Fe3+-K61E-1HGA | Fe2+-K61E-1HGA | Fe3+-K61N-1HGA | Fe2+-K61N-1HGA | Fe3+-H58Y-1HGA | Fe2+-H58Y-1HGA | Fe3+-H87Y-1HGA | Fe2+-H87Y-1HGA | Fe3+-H58L-1HGA | Fe2+-H58L-1HGA | Fe3+-H58L-3QJD | Fe2+-H58L-3QJD |
|---|---|---|---|---|---|---|---|---|---|---|---|---|---|---|---|---|
| Fe3+-1HGA | NA! | 0.0000 | 0.0000 | 0.0026 | 0.0000 | 0.0000 | 0.0001 | 0.0000 | 0.0001 | 0.0068 | 0.0000 | 0.0000 | 0.0000 | 0.0027 | 0.0018 | 0.3899 |
| Fe2+-1HGA | 0.0000 | NA! | 0.0000 | 0.0000 | 0.0000 | 0.1198 | 0.0052 | 0.0000 | 0.0000 | 0.0009 | 0.0000 | 0.0000 | 0.1241 | 0.0000 | 0.4945 | 0.0004 |
| Fe3+-H58R-1HGA | 0.0000 | 0.0000 | NA! | 0.0001 | 0.0000 | 0.0000 | 0.0000 | 0.0000 | 0.0005 | 0.0000 | 0.0000 | 0.0000 | 0.0000 | 0.0006 | 0.0000 | 0.0000 |
| Fe2+-H58R-1HGA | 0.0026 | 0.0000 | 0.0001 | NA! | 0.0000 | 0.0000 | 0.0000 | 0.0000 | 0.0687 | 0.0000 | 0.0000 | 0.0000 | 0.0000 | 0.3813 | 0.0001 | 0.0289 |
| Fe3+-K61E-1HGA | 0.0000 | 0.0000 | 0.0000 | 0.0000 | NA! | 0.0000 | 0.0000 | 0.0047 | 0.0000 | 0.0000 | 0.0000 | 0.0000 | 0.0000 | 0.0000 | 0.0267 | 0.0000 |
| Fe2+-K61E-1HGA | 0.0000 | 0.1198 | 0.0000 | 0.0000 | 0.0000 | NA! | 0.0882 | 0.0000 | 0.0000 | 0.0196 | 0.0000 | 0.0000 | 0.4695 | 0.0000 | 0.1979 | 0.0025 |
| Fe3+-K61N-1HGA | 0.0001 | 0.0052 | 0.0000 | 0.0000 | 0.0000 | 0.0882 | NA! | 0.0000 | 0.0000 | 0.1883 | 0.0000 | 0.0000 | 0.0899 | 0.0000 | 0.0883 | 0.0171 |
| Fe2+-K61N-1HGA | 0.0000 | 0.0000 | 0.0000 | 0.0000 | 0.0047 | 0.0000 | 0.0000 | NA! | 0.0000 | 0.0000 | 0.0000 | 0.0000 | 0.0000 | 0.0000 | 0.0089 | 0.0000 |
| Fe3+-H58Y-1HGA | 0.0001 | 0.0000 | 0.0005 | 0.0687 | 0.0000 | 0.0000 | 0.0000 | 0.0000 | NA! | 0.0000 | 0.0000 | 0.0000 | 0.0000 | 0.1134 | 0.0000 | 0.0007 |
| Fe2+-H58Y-1HGA | 0.0068 | 0.0009 | 0.0000 | 0.0000 | 0.0000 | 0.0196 | 0.1883 | 0.0000 | 0.0000 | NA! | 0.0000 | 0.0000 | 0.0606 | 0.0004 | 0.0525 | 0.1022 |
| Fe3+-H87Y-1HGA | 0.0000 | 0.0000 | 0.0000 | 0.0000 | 0.0000 | 0.0000 | 0.0000 | 0.0000 | 0.0000 | 0.0000 | NA! | 0.0007 | 0.0000 | 0.0000 | 0.0000 | 0.0000 |
| Fe2+-H87Y-1HGA | 0.0000 | 0.0000 | 0.0000 | 0.0000 | 0.0000 | 0.0000 | 0.0000 | 0.0000 | 0.0000 | 0.0000 | 0.0007 | NA! | 0.0000 | 0.0000 | 0.0000 | 0.0000 |
| Fe3+-H58L-1HGA | 0.0000 | 0.1241 | 0.0000 | 0.0000 | 0.0000 | 0.4695 | 0.0899 | 0.0000 | 0.0000 | 0.0606 | 0.0000 | 0.0000 | NA! | 0.0000 | 0.1808 | 0.0031 |
| Fe2+-H58L-1HGA | 0.0027 | 0.0000 | 0.0006 | 0.3813 | 0.0000 | 0.0000 | 0.0000 | 0.0000 | 0.1134 | 0.0004 | 0.0000 | 0.0000 | 0.0000 | NA! | 0.0000 | 0.0077 |
| Fe3+-H58L-3QJD | 0.0018 | 0.4945 | 0.0000 | 0.0001 | 0.0267 | 0.1979 | 0.0883 | 0.0089 | 0.0000 | 0.0525 | 0.0000 | 0.0000 | 0.0000 | 0.0000 | NA! | 0.0025 |
| Fe2+-H58L-3QJD | 0.0000 | 0.0000 | 0.0000 | 0.0000 | 0.0000 | 0.0000 | 0.0000 | 0.0000 | 0.0000 | 0.0000 | 0.0000 | 0.0000 | 0.0000 | NA! | 0.0000 | NA! |

*Table S2. Average, minimum, maximum and standard deviation for residue-wise RMSF values for the oxidized (Fe3+) and reduced (Fe2+) states of different Hb variants studied in this work. Four residues from each terminal have been excluded in this analysis since they are known to generally show larger RMSF value.*

| | Fe3+-1HGA | Fe2+-1HGA | Fe3+-H58R-1HGA | Fe2+-H58R-1HGA | Fe3+-K61E-1HGA | Fe2+-K61E-1HGA | Fe3+-K61N-1HGA | Fe2+-K61N-1HGA | Fe3+-H58Y-1HGA | Fe2+-H58Y-1HGA | Fe3+-H87Y-1HGA | Fe2+-H87Y-1HGA | Fe3+-H58L-1HGA | Fe2+-H58L-1HGA | Fe3+-H58L-3QJD | Fe2+-H58L-3QJD |
|---|---|---|---|---|---|---|---|---|---|---|---|---|---|---|---|---|
| average | 0.5817 | 0.5586 | 0.6382 | 0.5979 | 0.5424 | 0.5642 | 0.5690 | 0.5270 | 0.6102 | 0.5731 | 0.8738 | 0.8111 | 0.5645 | 0.6004 | 0.5584 | 0.5836 |
| min | 0.3790 | 0.3672 | 0.3820 | 0.3808 | 0.3608 | 0.3677 | 0.3703 | 0.3534 | 0.3906 | 0.3776 | 0.4081 | 0.4184 | 0.3634 | 0.3701 | 0.3478 | 0.3615 |
| max | 1.0727 | 1.0541 | 1.5795 | 1.2821 | 1.0776 | 1.1794 | 1.0371 | 1.1396 | 1.4744 | 1.1242 | 2.7310 | 2.4706 | 1.1589 | 1.4606 | 1.5834 | 1.1166 |
| stdev | 0.1664 | 0.1550 | 0.2558 | 0.1868 | 0.1568 | 0.1732 | 0.1544 | 0.1428 | 0.2101 | 0.1675 | 0.4433 | 0.3301 | 0.1693 | 0.1943 | 0.1998 | 0.1855 |

*Table S3. The average vertical ionization energies (<ΔE>, eV) for the redox states, total reorganization free energies (λ, eV), scaled λ values (scaling factor 1.6),[31] free energy change for the oxidation (ΔG°, eV), redox potential (E°, V), and selected α chain Hb mutants. The ET parameters for the Cyt b5:Hb complex (wild type) are also shown.*

| **Protein** | **Mutation** | **Avg. ΔE(Oxd)** | **Avg. ΔE(Red)** | **λ** | **Scaled λ** | **ΔG°** | ***E°*** | **Hb M formation** |
|---|---|---|---|---|---|---|---|---|
| Hb | Wild | 0.91 ± 0.007 | 3.10 ± 0.005 | 1.0965 ± 0.000 | 0.6853 | 2.004 ± 0.006 | -0.165[a] | < 1 % |
| α analogue of the Hb Zurich β variant | H58R | 0.95 ± 0.014 | 3.26 ± 0.002 | 1.1545 ± 0.0060 | 0.7216 | 2.105 ± 0.008 | -0.064 | NA |
| Hb Miyagi | K61E | 0.71 ± 0.014 | 3.04 ± 0.011 | 1.1675 ± 0.0018 | 0.7297 | 1.873 ± 0.013 | -0.296 | Increased Hb M formation |
| Hb J-Buda | K61N | 0.89 ± 0.009 | 3.04 ± 0.001 | 1.0707 ± 0.0041 | 0.6692 | 1.965 ± 0.005 | -0.204 | NA |
| Hb Kirklareli | H58L | 0.87 ± 0.102 | 3.11 ± 0.005 | 1.1201 ± 0.0488 | 0.7001 | 1.993 ± 0.054 | -0.176 | Increased Hb M formation |
| Hb Kirklareli (3QJD) | H58L | 0.92 ± 0.001 | 3.12 ± 0.002 | 1.0987 ± 0.0008 | 0.6867 | 2.019 ± 0.001 | -0.150 | |

| | | | | | | | | |
|---|---|---|---|---|---|---|---|---|
| Cyt b5 (3NER) | Wild | 6.13 ± 0.010 | 9.10 ± 0.004 | 1.4878 ± 0.0031 | 0.9299 | 7.615 ± 0.007 | -0.212[a] | -- |
| Cyt b5:Hb complex (3NER:1HGA) | Wild | 3.70 ± 0.021 | 8.13 ± 0.046 | 2.2145 ± 0.0331 | 1.381 | 5.913 ± 0.012 | -0.047[b] | -- |

[a] Computed redox potential (E°) shifted to match experimental E° versus normal hydrogen electrode. Same shift constant used for all mutants.
[b] Experimental driving force for ET from Cyt b5 to Hb.
NA Information not available in the Hbvar database and literature.

*Table S4. The average vertical ionization energies (<ΔE>) for the protein redox states (solvent molecules were stripped before single point energy estimation with sander), the protein and total reorganization free energies ($\lambda_{prot}$, and λ), solvent reorganization energy ($\lambda_{solv}$) for Hb wild and selected mutants.*

| Hb α chain Variants | Mutation | Protein Avg. ΔE(Oxd) | Protein Avg. ΔE(Red) | ΔG° | λ | $\lambda_{prot}$ | $\lambda_{solv} = (\lambda - \lambda_{prot})$ | % contribution of $\lambda_{solv}$ to λ |
|---|---|---|---|---|---|---|---|---|
| Hb | Wild | -0.83 | -0.32 | 2.00 | 1.0965 | 0.26 | 0.84 | 76.43 |
| Hb M Boston | H58Y | -0.88 | -0.33 | 2.00 | 1.1094 | 0.28 | 0.83 | 75.02 |
| α analogue of the Hb Zurich β variant | H58R | 0.29 | 0.86 | 2.10 | 1.1545 | 0.29 | 0.87 | 75.21 |
| Hb Miyagi | K61E | -3.02 | -2.95 | 1.87 | 1.1675 | 0.04 | 1.13 | 96.81 |
| Hb J-Buda | K61N | -1.71 | -1.60 | 1.97 | 1.0707 | 0.05 | 1.02 | 95.28 |
| Hb Kirklareli | H58L | -0.82 | -0.23 | 1.99 | 1.1201 | 0.30 | 0.82 | 73.38 |

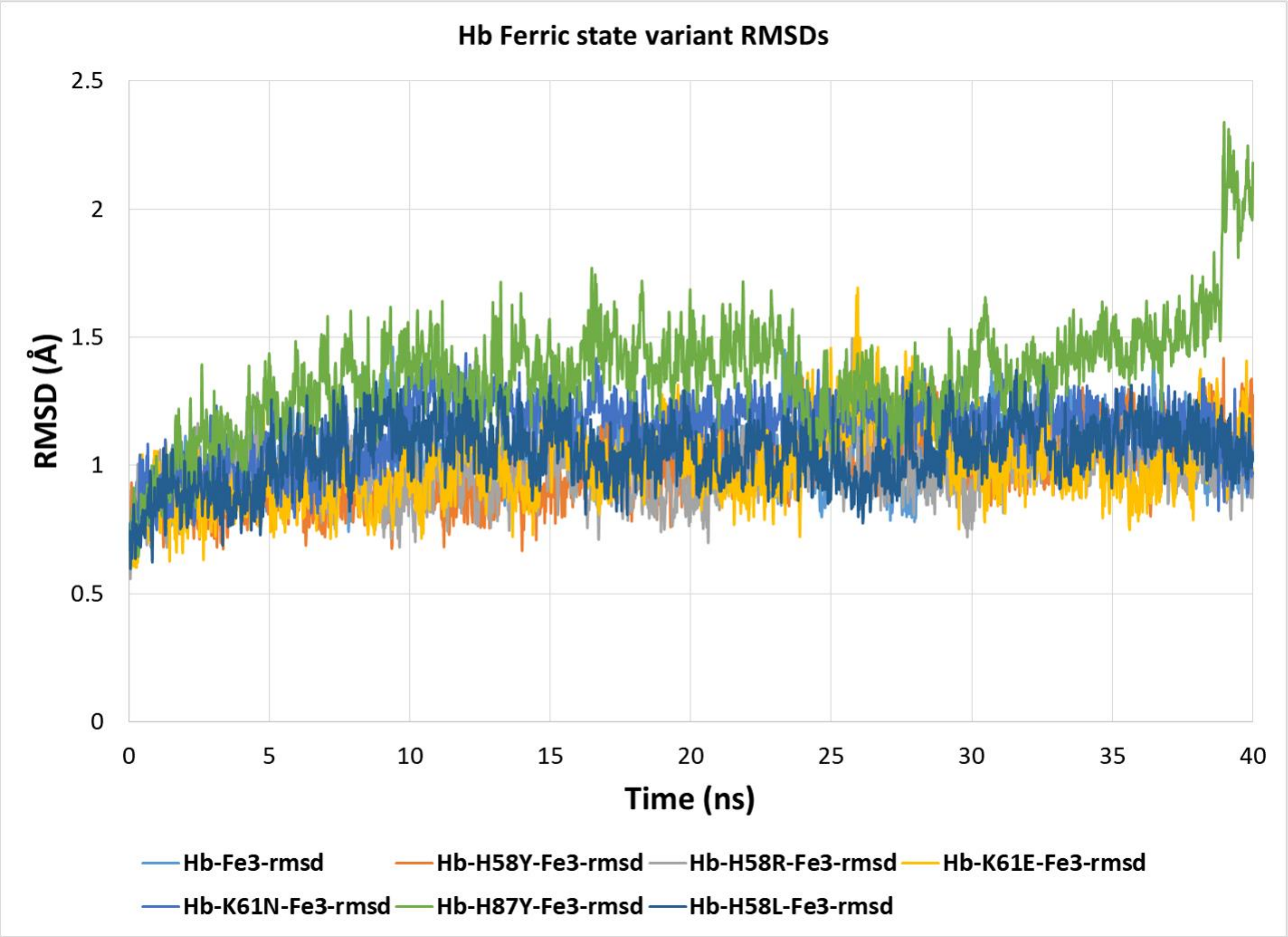

*Figure S4. RMSD of the Hb A alpha chain and its variants in the Fe2+ state during the 40 ns MD simulation.*

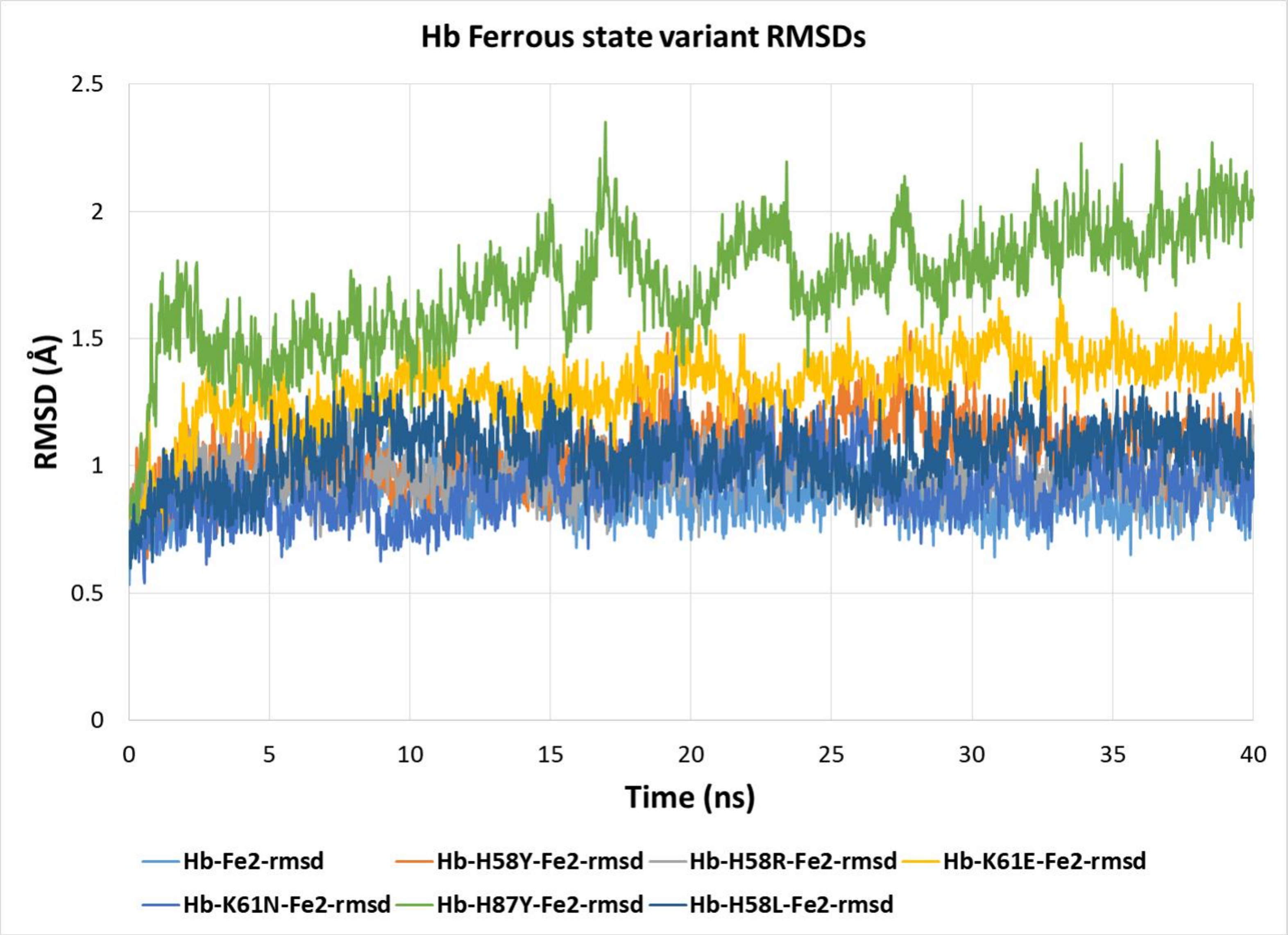

*Figure S5. RMSD of the Hb A alpha chain and its variants in the Fe3+ state during the 40 ns MD simulation.*

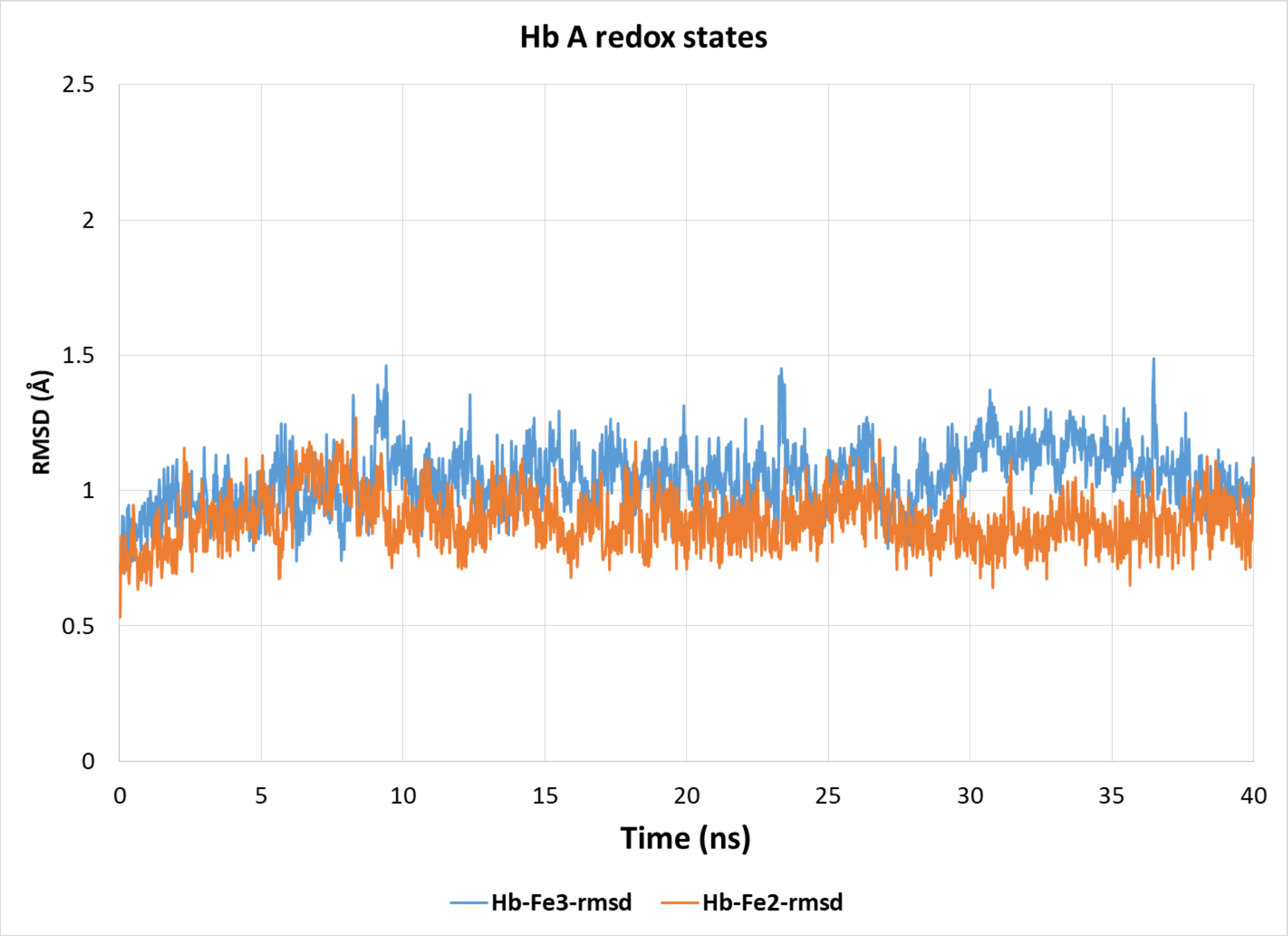

*Figure S6. RMSD of the Hb A alpha chain redox states during the 40 ns MD simulation.*

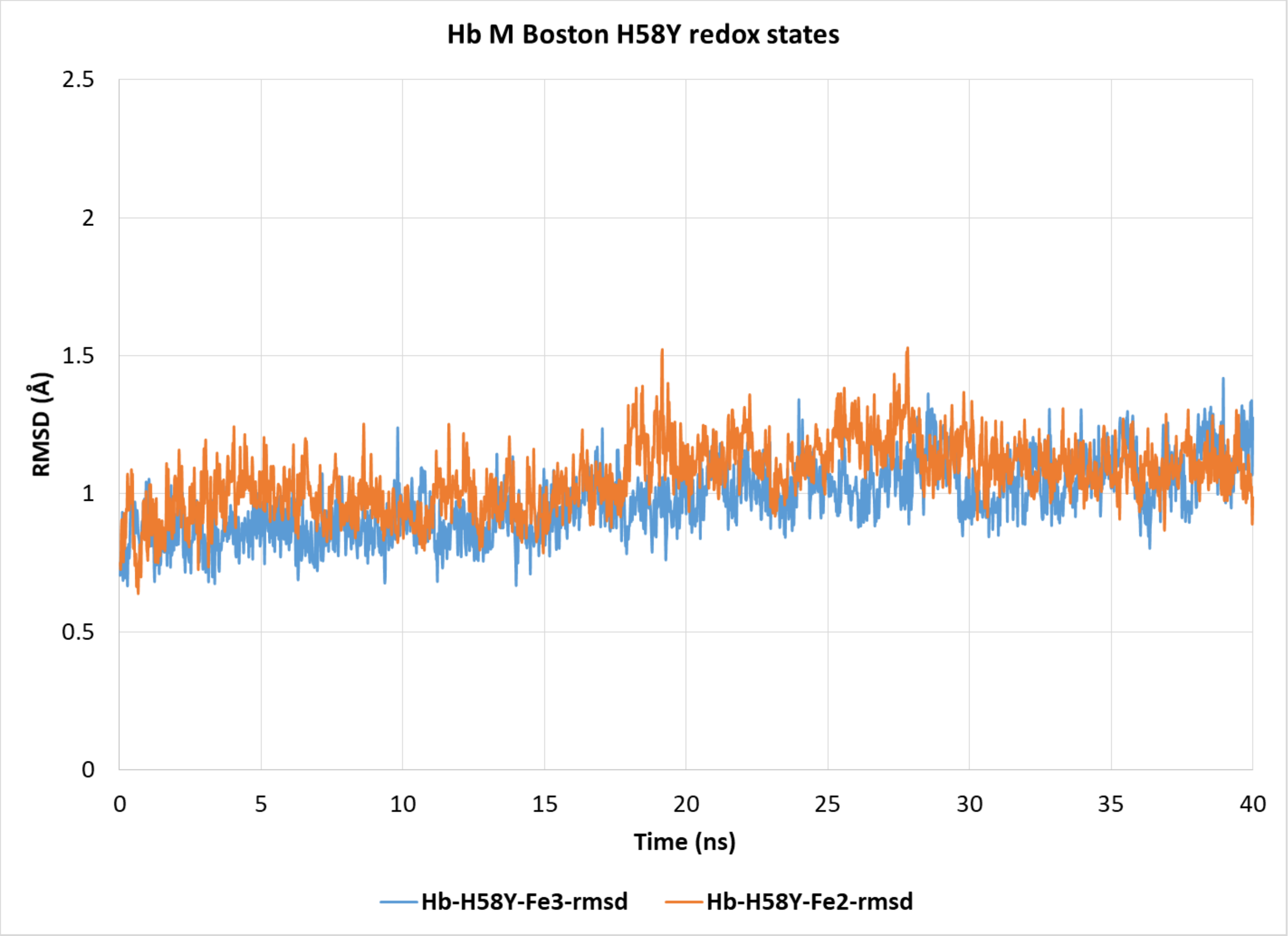

*Figure S7. RMSD of the Hb M Boston alpha chain redox states during the 40 ns MD simulation.*

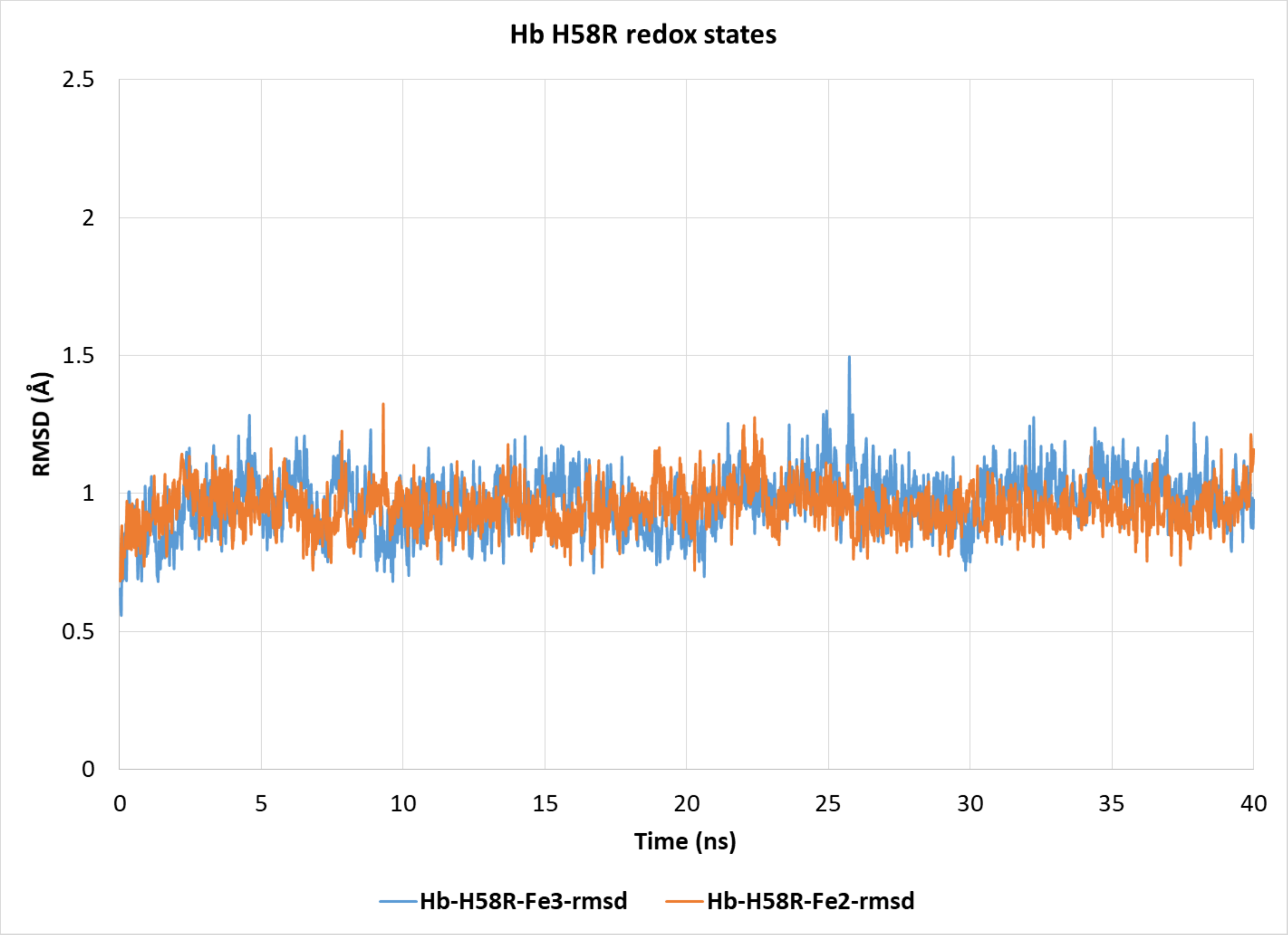

*Figure S8. RMSD of the Hb H58R variant alpha chain redox states during the 40 ns MD simulation.*

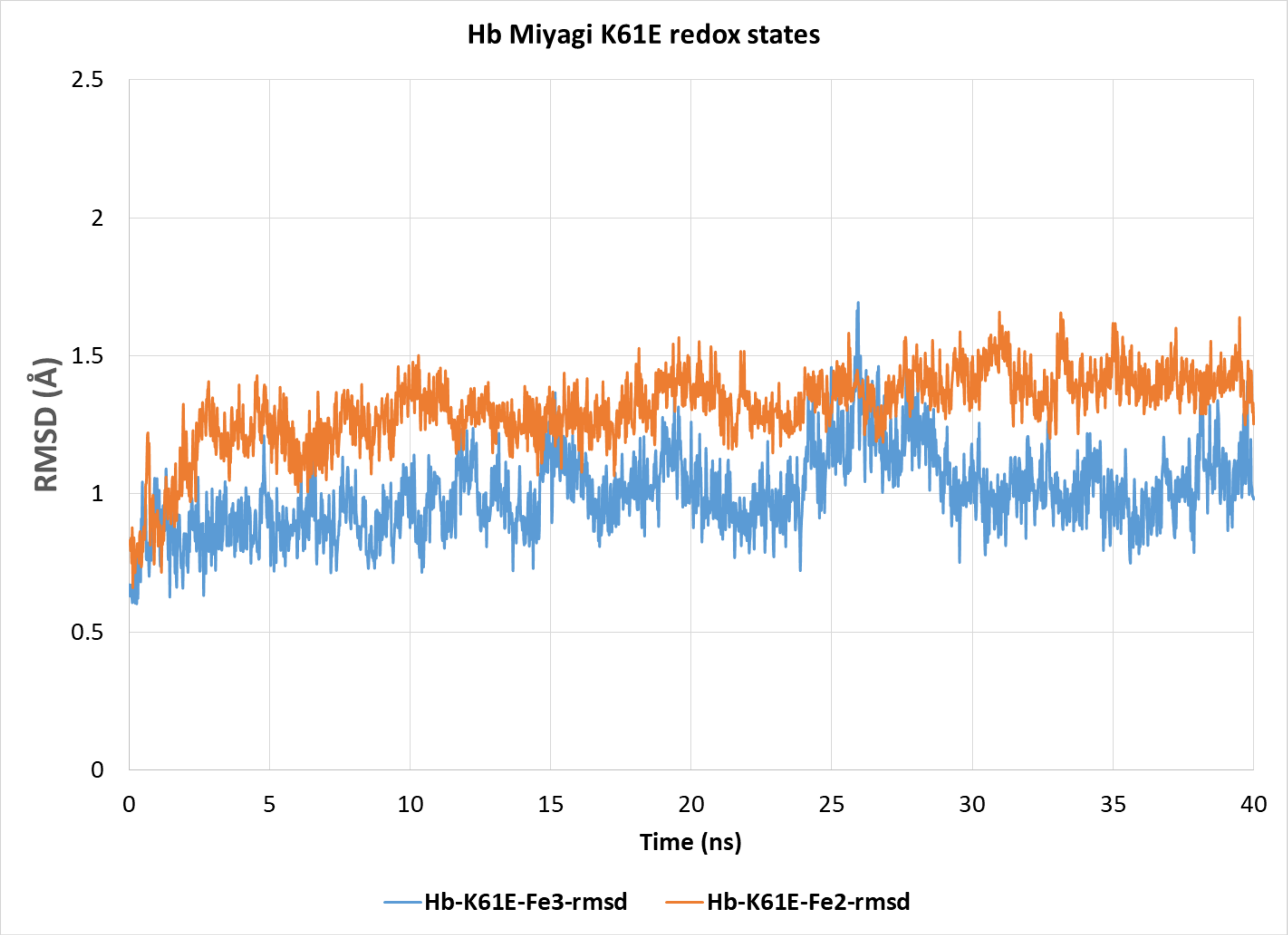

*Figure S9. RMSD of the Hb Miyagi variant alpha chain redox states during the 40 ns MD simulation.*

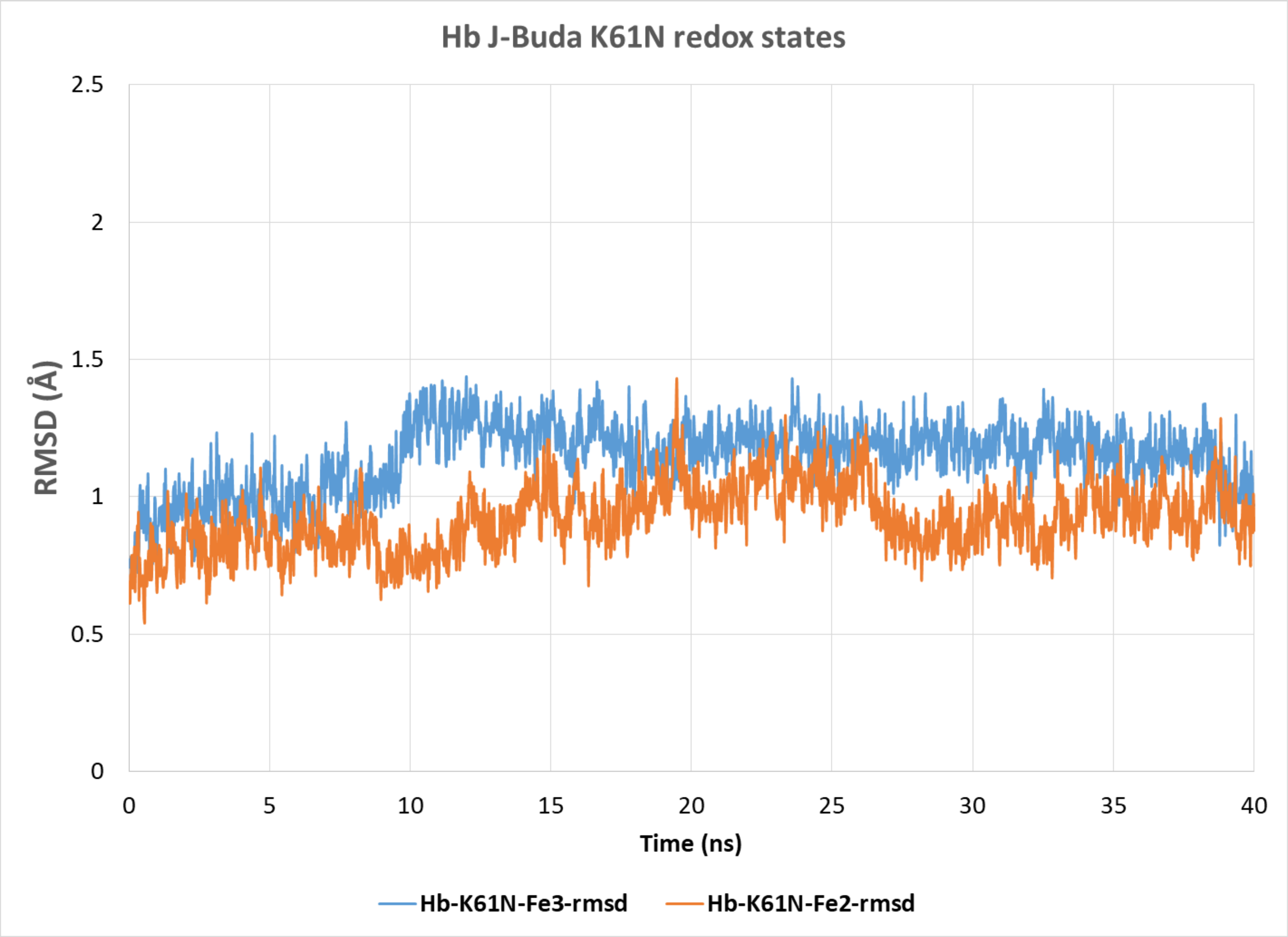

*Figure S10. RMSD of the Hb J-Buda variant alpha chain redox states during the 40 ns MD simulation.*

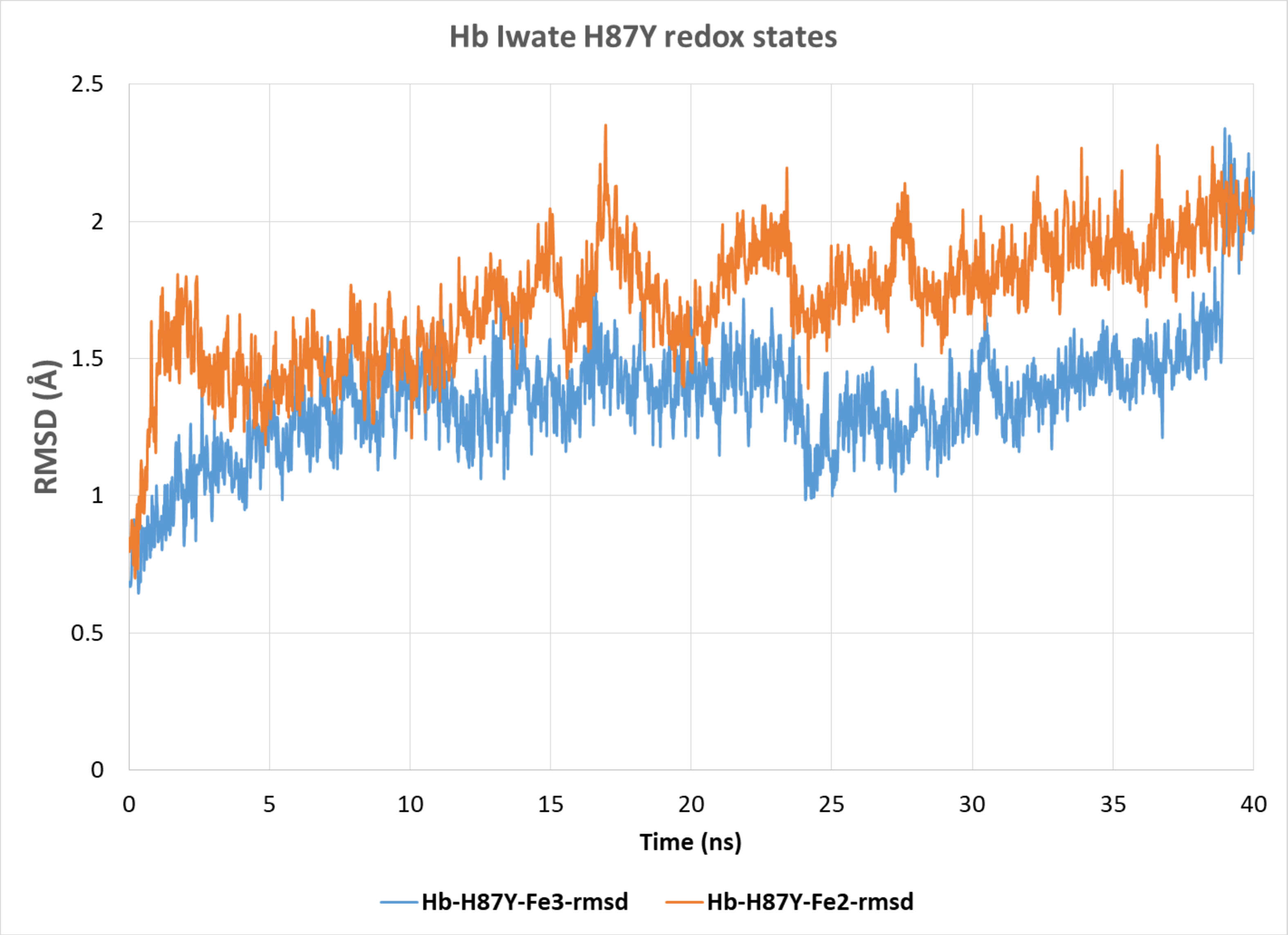

*Figure S11. RMSD of the Hb Iwate variant alpha chain redox states during the 40 ns MD simulation.*

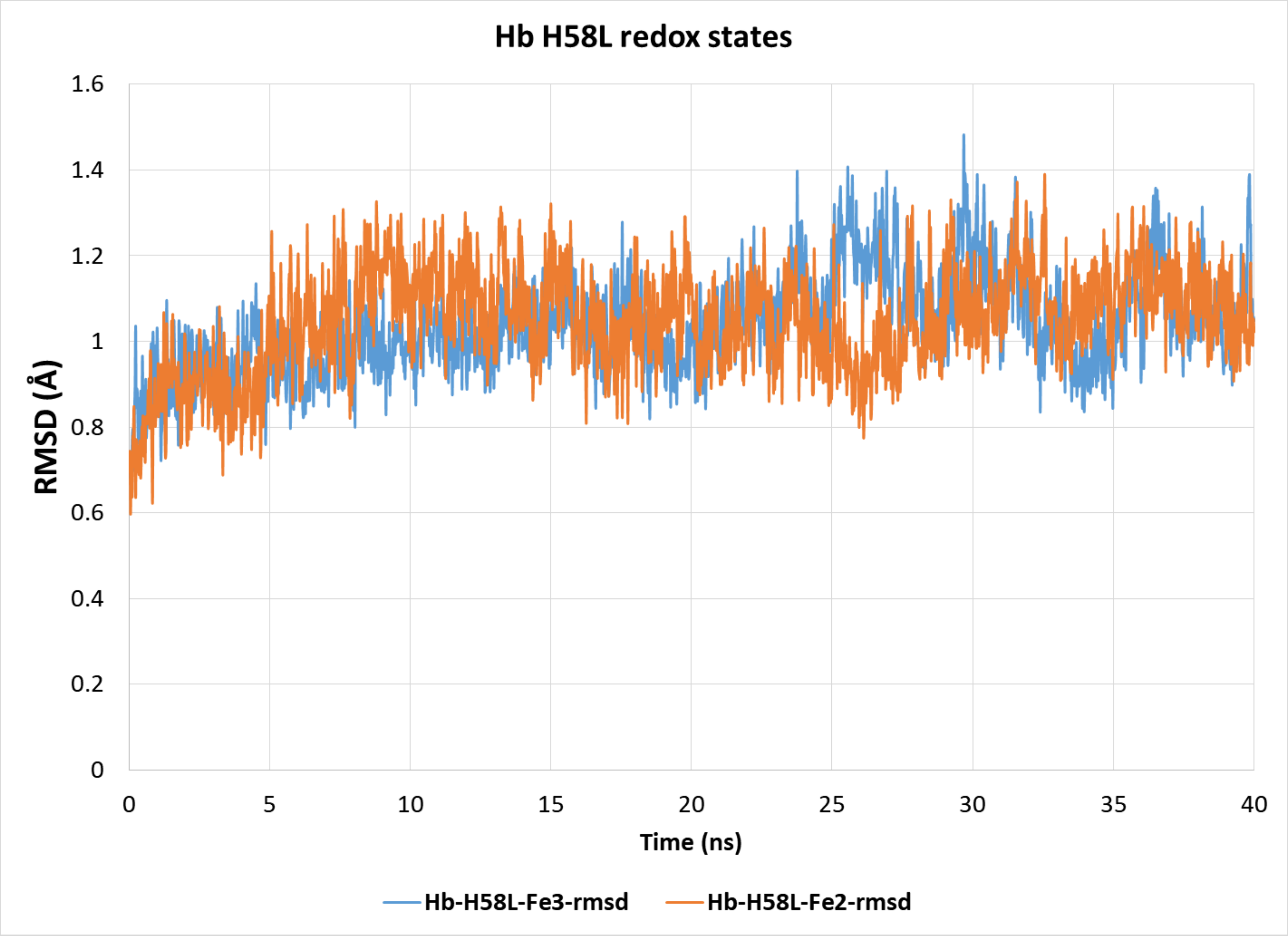

*Figure 12.. Figure S13. RMSD of the Hb Kirklareli variant alpha chain redox states during the 40 ns MD simulation.*
